# Joint species distributions reveal the combined effects of host plants, abiotic factors and species competition as drivers of species abundances in fruit flies

**DOI:** 10.1101/2020.12.07.414326

**Authors:** Benoit Facon, Abir Hafsi, Maud Charlery de la Masselière, Stéphane Robin, François Massol, Maxime Dubart, Julien Chiquet, Enric Frago, Frédéric Chiroleu, Pierre-François Duyck, Virginie Ravigné

**Author notes:** **Cite as:** Facon B, Hafsi A, Charlery de la Masselière M, Robin S, Massol F, Dubart M, Chiquet J, Frago E, Chiroleu F, Duyck P-F, Ravigné V (2021) Joint species distributions reveal the combined effects of host plants, abiotic factors and species competition as drivers of community structure in fruit flies. bioRxiv, 2020.12.07.414326. ver. 4 peer-reviewed and recommended by Peer community in Ecology. https://doi.org/10.1101/2020.12.07.414326.

## Abstract

The relative importance of ecological factors and species interactions for phytophagous insect species distributions has long been a controversial issue. Using field abundances of eight sympatric Tephritid fruit flies on 21 host plants, we inferred flies’ realized niches using joint species distribution modelling and network inference, on the community as a whole and separately on three groups of host plants. These inferences were then confronted to flies’ fundamental niches estimated through laboratory-measured fitnesses on host plants. Species abundances were mainly determined by host plants followed by climatic factors, with a minor role for competition between species sharing host plants. The relative importance of these factors mildly changed when we focused on particular host plant groups. Despite overlapping fundamental niches, specialists and generalists had almost distinct realized niches, with possible competitive exclusion of generalists by specialists on Cucurbitaceae, and different assembly rules: specialists were mainly influenced by their adaptation to host plants while generalist abundances varied regardless of their fundamental host use.

## Introduction

The search for fundamental processes underlying species distributions is among the oldest challenges in ecology (Diamond 1975, Gotelli and Graves 1996). Understanding assembly processes could also be crucial for coping with global changes and habitat loss currently affecting both abiotic conditions and species distributions (Adler and HilleRisLambers 2008). Species distributions can be determined by several factors such as environmental filtering, interspecific interactions, regional species pool and dispersal (Müller *et al*. 2011, D’Amen *et al*. 2018, Nakadai *et al*. 2018, Jabot *et al*. 2020). Despite decades of research, estimating the relative importance of these processes on species distributions has proven particularly complex (Pollock *et al*. 2014, Zurell *et al*. 2018). Most of these processes imprint species distributions in a scale-dependent manner (Meynard *et al*. 2013). For instance, abiotic factors are generally thought to determine large-scale species ranges, whereas interspecific interactions would influence species distributions at smaller spatial scales (Heikkinen *et al*. 2007, Thuiller *et al*. 2015, but see Gotelli et al 2010 and Araújo and Rozenfeld 2014).

Phytophagous insects are among the most diverse and abundant groups of terrestrial animals and a major component of ecosystems due to their tight interaction with primary producers, and their sometimes important economic impacts and invasive potential (Roy *et al*. 2015). Knowledge of the main determinants of insect occurrence on particular plant species and their potential to colonize and persist in a given area is however still limited. In particular, the importance of interspecific competition in structuring phytophagous insect communities has been a controversial issue (Kaplan and Denno 2007). Many experimental studies conclude that interspecific competition plays a primary role (Denno *et al*. 1995, Kaplan & Denno 2007), but the consistent absence of negative co-occurrence patterns in natural phytophagous insect communities suggests otherwise (Tack *et al*. 2009, Brazeau & Schamp 2019). This apparent discrepancy could result from the regulation of phytophagous insect populations below competitive levels through shared predators or parasites (Hairston *et al*. 1960). Phytophagous insects would also rarely overexploit their hosts, leaving sufficient plant material for competition with other species to be mild (Kaplan & Denno 2007). Ecological differences between species could lower the intensity of competition (Stewart *et al*. 2015). Lastly, some phytophagous arthropods could benefit from previous attack of the host plant by other species (Godinho *et al*. 2016). The importance of competition relative to ecological conditions in shaping phytophagous insect distributions thus remains an open question and demands appropriate testing (Augustyn *et al*. 2016, Nakadai *et al*. 2018).

Whether biotic interactions affect species distributions should be uncovered from proper analysis of patterns of co-occurrence. Species interactions are expected to affect species occurrence, *e*.*g*., competition should cause checkerboard patterns of occurrence. However, species occurrences also result from common or diverging species dependence on confounding environmental factors. Species that share the same abiotic niche will frequently co-occur without necessarily interacting (Wisz *et al*. 2013, Blanchet *et al*. 2020). Conversely, negative co-occurrence patterns may simply result from diverging ecological requirements. As a consequence, estimating the effect of species interactions on species distributions first requires properly characterizing species abundance’s responses to environmental variables (Pollock *et al*. 2014), which is the object of species distribution modeling (SDM) approaches (Elith & Leathwick 2009). A particular class of SDM approaches, joint species distribution models (JSDM) attempts to infer the relationships between species abundances and environmental variables, explicitly accounting for the interdependence of species distributions using multivariate regression methods (Pollock *et al*. 2014). In addition to estimates of the effect of habitat filtering on species distributions, these approaches provide residual covariances between species abundances, *i*.*e*., covariances not explained by environmental factors. Residual covariances result from species interactions and a diversity of other factors, as *e*.*g*., missing covariates, so that there is no simple relationship between species interactions and residual covariances (Zurell *et al*. 2018).

To further track species interactions, a growing body of literature pleads for using independent knowledge of species traits in species distribution modelling (Lavorel *et al*. 1997, Kraft *et al*. 2008, Poisot *et al*. 2015), which is still seldom done. Explicitly comparing estimates of fundamental niche, *i*.*e*., measures of fitness in controlled conditions and in absence of species interactions, to estimates of realized niche, *i*.*e*., inferred species abundances’ responses to environmental factors, could shed new light on the gap between fundamental and realized niches and the importance of species interactions in shaping species distributions.

In the case of phytophagous insects, an obvious feature of the environment to account for is host plant identity. Host plants can be treated as any environmental cofactor, and their effects on species abundances can be inferred directly from adequate abundance data (Ferrier and Guisan 2006). Host plants impose a specific challenge because modelling a phytophagous community as a whole relies on the assumption that the interdependence of species abundance does not depend on host plants. But as intraguild interactions mostly occur in/on plant organs, they may be modulated by plant species identity, with possible consequences for species occurrence patterns (Ulrich *et al*. 2017). Analyzing competition patterns on different host plants could therefore allow detecting the role of host plants in shaping species co-occurrences.

Here we aimed at disentangling the roles of host plant species, abiotic factors, and interspecific interactions on the distributions of eight fruit fly species (Tephritidae) occurring in sympatry on a diversity of host plants and in highly variable abiotic conditions. The study system, which comprises four generalist species, three specialists of Cucurbitaceae, and one specialist of Solanaceae, presents key advantages to tackle community assembly questions. First, these eight species occupy a small island in South-western Indian Ocean (Réunion, 2512 km2) where they are considered the main actors in the guild of fruit-eating phytophagous arthropods (Quilici and Jeuffrault, 2001). Second, the local environment is characterized by important variability in elevation (from 0 to 3000m), climatic conditions, land use, and plant distributions (Duyck *et al*. 2006a). Observational and experimental studies have suggested that climatic factors could influence local Tephritid distributions (Duyck *et al*. 2004). Climatic factors were even found more influential than host plant diversity in allowing coexistence in an analysis of the distributions of the four generalist species on four host plants (Duyck *et al*. 2008). Lastly, competition between the eight species has repeatedly been advocated to shape this community. First, host-use strategies largely overlap, opening possibilities of competition (Quilici & Jeuffrault 2001, Duyck *et al*. 2008). Second, the arrival of one generalist species on the island has constrained the host ranges of some resident species, without complete exclusion (Charlery de la Masselière *et al*. 2017a). Third, larval competition experiments in a subset of plant species and abiotic conditions have evidenced hierarchical competition interactions among the generalist species (Duyck *et al*. 2006b).

Here we confronted a long-term field dataset describing abundances of the eight fly species on 21 host plants with laboratory measures of fundamental host use obtained for seven of the fly species on the same plants. We first modelled joint species distributions using Poisson-LogNormal (PLN) modelling (Chiquet *et al*. 2019) and conducted model selection among various combinations of host plant species and ecological covariates (representing temperature, rainfall, elevation, land use and date). Residual correlations estimated under the selected model were further dissected to identify significant traces of unexplained co-occurences. Second, we assessed whether knowledge on fundamental host use was sufficient to explain field species abundances by accounting for host plant species either directly or through estimates of female preference and larval performance in laboratory conditions. Finally, we tested for a potential dependence of community structuring factors on host plants by replicating the analyses on three subsets of plants: Cucurbitaceae, Solanaceae and the other plants.

## Methods

### Species abundance table

Field campaigns were conducted over a period of 18 years (1991-2009) to identify potential host plants for Tephritidae on the whole Réunion island including orchards, gardens, and natural areas. These surveys were assembled in a previous study (Charlery de la Masselière *et al*. 2017a), and used here as species abundance table. Each observation corresponds to the number of individual flies of the eight species recorded from a set of fruits sampled in one location at a specific date. For each sample, the collected fruits were counted and weighted, before being stored until adult fly emergence. To avoid keeping samples that could have suffered from a transportation or storage issue, only samples with at least one individual fly were kept. Among these, we further selected samples with GPS coordinates, and belonging to one of the 21 host plants characterized in the laboratory (see below). Of the 12872 initial samples, we therefore kept 4918 samples, and a total of 97351 individual flies. Samples covered 104 field sessions all year round over the study period (Tables S1 and S2) and originated from 380 sites well distributed over the whole island (Figure 1A). Additional details on sample collection can be found in Appendix S1.

**Figure 1:**
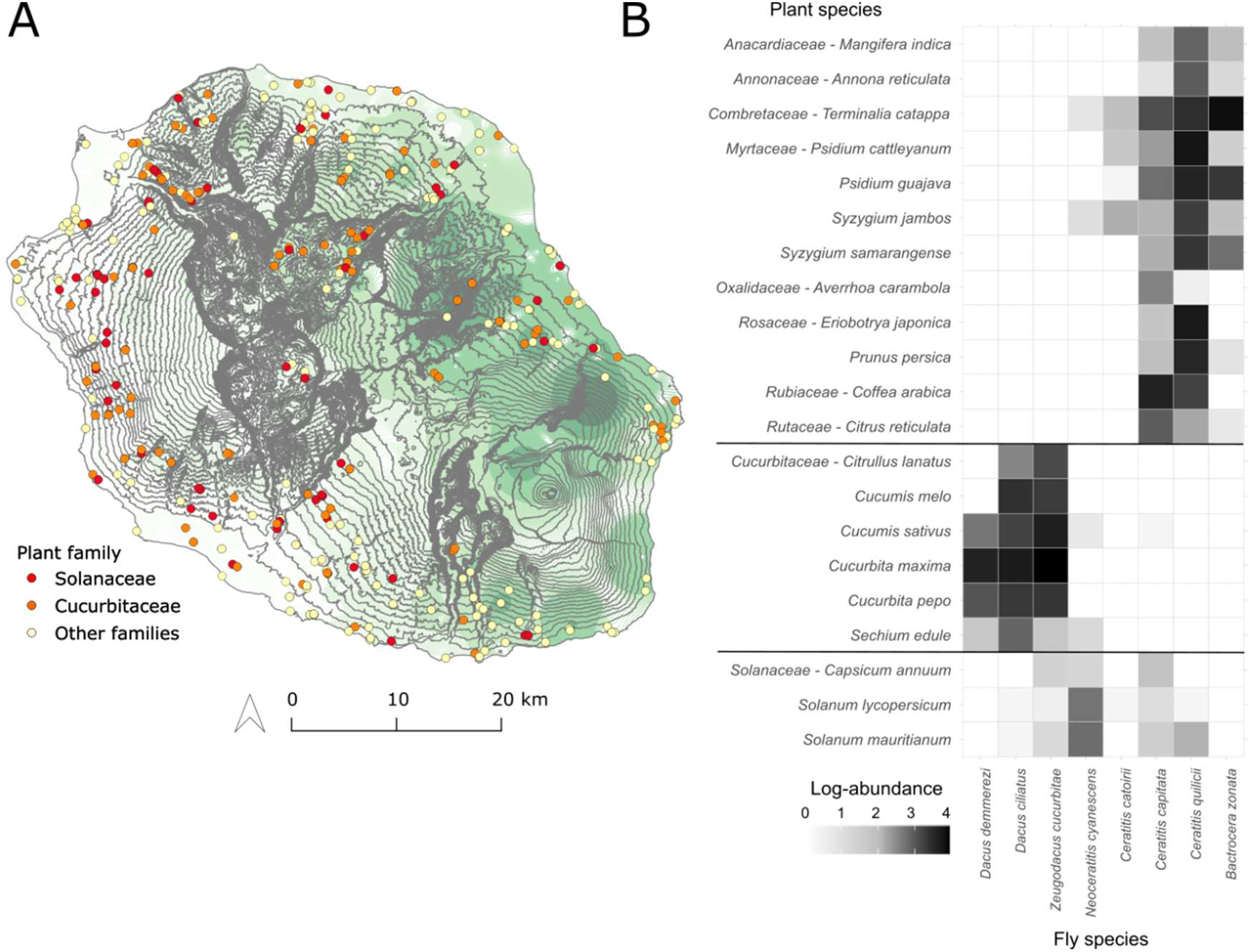
Characteristics of the species abundance dataset. A) Sampling sites in Réunion. Dot colors refer to the family of the sampled plant (Solanaceae n=259, Cucurbitaceae n=2347, other families n=2285 samples). Elevation, represented by grey isoclines, ranges from 0 along the coast, to >3000 m, in the center of the island, and strongly correlates with annual temperature. Pluviometry (here median rainfall over 1986-2016) is represented by green isohyets. The island is separated into two contrasted rainfall regimes: very humid all year round in the east, and drier, especially during winter, in the west. B) Fly species abundances on the 21 studied plant species.

### Ecological covariables

The GPS coordinates of each sample were used to retrieve ecological and climatic characteristics from GIS information available on the CIRAD Agricultural Web Atlas for Research (AWARE, https://aware.cirad.fr). Each sample was associated with a month and a year, a land use category, an elevation, three pluviometry descriptors (minimal rainfall in the 20% most humid years, minimal rainfall in the 20% driest years, median annual rainfall between 1986 and 2016), and three temperature variables (minimal, mean, maximal annual temperature between 1987 and 2017) (Figure 1A, Appendix S1). To account for correlations between some of the variables, a FAMD (factorial analysis of mixed data) was conducted on all 10 variables using *FactoMineR* (Lê *et al*. 2008). Ten uncorrelated dimensions were obtained and subsequently used as ecological covariates in the following analyses (Figure S2 for details on the FAMD).

### Species traits

For all species but *Dacus ciliatus*, fundamental host use, *i*.*e*., fly fitness on host plants in optimal abiotic conditions and in absence of antagonists, was characterized using four traits describing larval performances and female preferences for 21 plant species. Female preferences were the numbers of eggs laid by females during 24h on each of the 21 fruit species in the ‘no-choice’ experiment of Charlery de la Masselière *et al*. (2017b). Larval performances (survival probability until maturity *s*, development time *T*, and pupal weight *w*) were obtained from Hafsi *et al*. (2016) for 17 plant species, and in the current study for *Coffea arabica, Solanum mauritianum, Syzygium jambos*, and *S. samarangense*, using the same methods. The three larval performance traits were combined into a single performance trait using the formula:

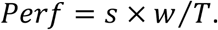

Both preference and performance traits were log-transformed before being included as covariates in statistical models.

### Statistical analysis

#### Datasets

To account for the possibility that determinants of species distributions depend on host plant identity, we replicated all analyses on the full 21-plants dataset, and on the following three sub-datasets: (i) Cucurbitaceae only (3 fly species x 6 plant species, 2347 samples), (ii) Solanaceae only (3 fly species x 3 plant species, 259 samples), (iii) other plant families (4 fly species x 12 plant species, 2285 samples).

#### Statistical modelling

Joint variations in fly species abundances were modelled using Poisson-LogNormal (PLN) models with the *PLN()* function in the *PLNmodels* R package (Chiquet *et al*. 2018, 2019). A PLN model is a multivariate mixed generalized linear model, where each species count is assumed to arise from a Poisson distribution with a parameter resulting from fixed effects of covariates and a random lognormal effect. Random effects associated to all species observed in a sample are jointly sampled from a multivariate lognormal distribution. The residual variance-covariance of the multivariate distribution reveals species abundance covariations still unexplained after controlling for confounding environmental covariates and differences in sampling efforts. Samples differed in fruit number and weight, inducing uncontrolled variation in sampling effort among samples potentially leading to spurious associations between fly species abundances. Consequently, the (log-transformed) total fruit weight of each sample was added as an offset to every tested model (fruit number was also tested as an offset yielding identical conclusions, results not shown). Model diagnostics were conducted using the R package *DHARMa* (Hartig 2020). Covariations between species abundances unexplained by covariates were further investigated using the *PLNnetwork()* function, which adjusts the considered model under a sparsity constraint on the inverse of the variance-covariance matrix, *i*.*e*., constraining the number of edges in the resulting estimated network. The stability of the resulting species associations was estimated as their selection frequency in bootstrap subsamples of the StARS model selection procedure (range 0-1; Liu *et al*. 2010; Appendix S3).

#### Model selection design

The importance of plant species identity, ecological covariates and species interactions to explain species abundances was approached by model selection using the extended BIC criteria (Chen & Chen 2008). First we focused on species response to environmental variables and compared models (listed in Table 1) including either no covariate (Model 1-0), plant species identity as a cofactor (Model 1-1), ecological variables (all 10 FAMD dimensions described above, Model 1-3) or both plant species and ecological variables (Model 1-5). Second, we evaluated the importance of residual species abundance covariations. *PLNmodels* enables fitting models where the residual variance-covariance matrix is constrained to be diagonal. Such models assume no possible interaction between fly species. We therefore compared all models with their diagonal counterpart (Models 1-2, 1-4 and 1-6). Third we estimated how well knowledge of the fundamental niche, here laboratory-measured preferences and performances, explained field abundances. In its present form, *PLNmodels* does not allow accounting for species traits (covariates describe samples but not species within samples). To cope with this limitation, we considered that assuming that species distribute according to their fundamental niche implies that species interactions are negligible, and species distribute independently from one another. Using this assumption, we fitted models separately for each fly species, obtained their likelihoods and numbers of parameters, and computed the BIC of the seven-species dataset as:

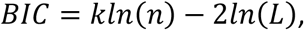

where *k* and *L* are the sums of the numbers of parameters and likelihoods over the seven single-species datasets and *n* the number of samples. Following this principle, we built models where the host plant cofactor was replaced by either preference (Model 2-7 without ecological covariates and 2-8 with ecological covariates), or performance (Models 2-9 and 2-10), or both preference and performance (Models 2-11 and 2-12). In addition, all previous models were reevaluated on the datasets excluding the species lacking fundamental host use estimates (*D. ciliatus*; Models 2-0 to 2-6).

**Table 1:** Model selection on the 21-plant dataset (*n* = 4,918). Models are ranked by increasing BIC (from best to worst). *k* is the number of parameters. *L* is the log-likelihood. Δ_BIC_ is the BIC difference between any focal model and the best one.

| Models | Covariates | Residual matrix | $K$ | $L$ | BIC | $\Delta_{\text{BIC}}$ |
| --- | --- | --- | --- | --- | --- | --- |
| A) Model set 1 (21 plants x 8 flies) |  |  |  |  |  |  |
| Model 1-5 | Plant + Eco | Full | 284 | -27664.3 | 57742.8 | 0.0 |
| Model 1-6 | Plant + Eco | Diagonal | 256 | -27997.8 | 58171.7 | 428.9 |
| Model 1-2 | Plant | Diagonal | 176 | -28608.9 | 58713.9 | 971.1 |
| Model 1-1 | Plant | Full | 204 | -28784.0 | 59302.1 | 1559.2 |
| Model 1-3 | Eco | Full | 124 | -35888.8 | 72831.6 | 15088.8 |
| Model 1-4 | Eco | Diagonal | 96 | -36598.4 | 74012.9 | 16270.1 |
| Model 1-0 | None | Full | 44 | -37228.1 | 74830.3 | 17087.4 |
| B) Model set 2 (21 plants x 7 flies) |  |  |  |  |  |  |
| Model 2-6 | Plant + Eco | Diagonal | 224 | -20753.9 | 43374.1 | 0.0 |
| Model 2-5 | Plant + Eco | Full | 245 | -20723.1 | 43487.5 | 113.4 |
| Model 2-2 | Plant | Diagonal | 154 | -21563.1 | 44409.4 | 1035.3 |
| Model 2-1 | Plant | Full | 175 | -21504.7 | 44467.5 | 1093.4 |
| Model 2-12 | Preference + Performance + Eco | Diagonal | 98 | -23924.4 | 48665.3 | 5291.2 |
| Model 2-10 | Performance + Eco | Diagonal | 91 | -24387.4 | 49533.0 | 6158.9 |
| Model 2-11 | Preference + Performance | Diagonal | 28 | -25689.9 | 51613.0 | 8238.9 |
| Model 2-8 | Preference + Eco | Diagonal | 91 | -25516.9 | 51792.1 | 8418.0 |
| Model 2-9 | Performance | Diagonal | 21 | -26443.3 | 53061.6 | 9687.5 |
| Model 2-4 | Eco | Diagonal | 84 | -27082.1 | 54864.0 | 11489.9 |
| Model 2-7 | Preference | Diagonal | 21 | -27628.9 | 55432.8 | 12058.7 |
| Model 2-3 | Eco | Full | 105 | -27321.5 | 55517.8 | 12143.7 |
| Model 2-0 | None | Full | 35 | -31060.1 | 62411.8 | 19037.7 |

## Results

The variance-covariance structure of the complete dataset (8 fly x 21 plant species) was first inferred by fitting a PLN model without any covariate (Model 1-0). The obtained residual variance-covariance matrix (Figure 2A) revealed a sharp distinction between three groups of flies: (i) the four generalists (*Bactrocera zonata, Ceratitis capitata, C. catoirii*, and *C. quilicii*), (ii) the three specialists of Cucurbitaceae (*D. ciliatus, D. demmerezi* and *Zeugodacus cucurbitae*), and the specialist of Solanaceae (*Neoceratitis cyanescens*). While *N. cyanescens* abundances showed very low covariances with other species, the two other groups showed positive within-group covariances and negative among-group covariances. This variance-covariance structure suggested strong separation of the realized niches of the three groups, likely mediated by host plants.

**Figure 2:**
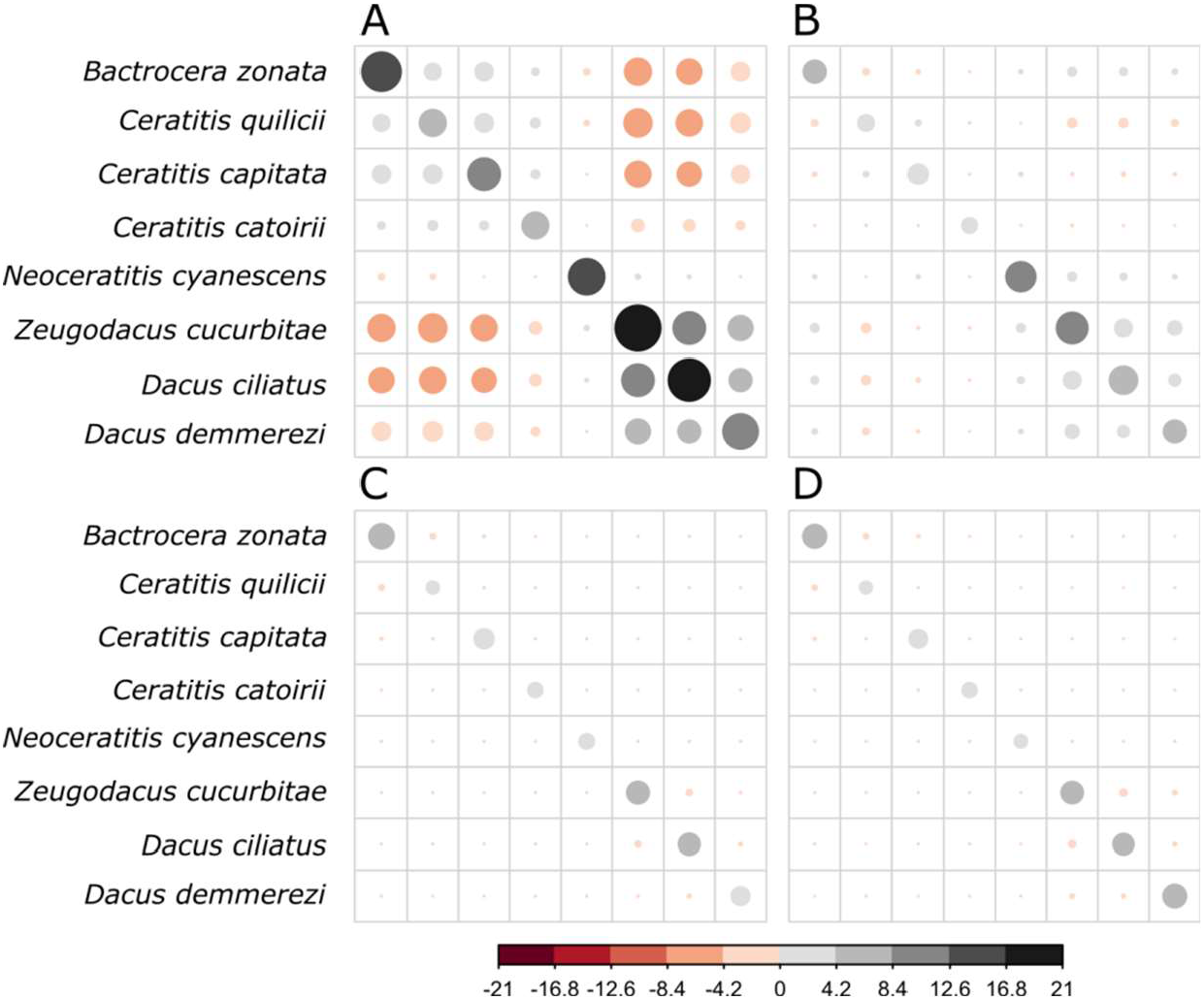
Residual variance-covariance matrices obtained after PLN model fitting on species abundances. A) Without any covariate (Model 1-0). B) With ecological covariates (Model 1-3). C) With plant species as a cofactor (Model 1-1). D) With both plant species and ecological covariates (Model 1-5).

Considering model selection between models without species traits, *i*.*e*., models including combinations of plant species and ecological covariates (Table 1), model ranking was equivalent on the eight-species (Models 1-0 to 1-6) and seven-species (Models 2-0 to 2-6) datasets, except that diagonal models tended to be slightly better than their full-matrix counterparts on the latter. This result was further shown robust to sample bootstrapping (Appendix S3).

For both datasets, the selected model included both ecological variables and host plant species as covariates (Models 1-5 and 2-6), which strongly improved the BIC (Δ_BIC_ = 17087.4 and 18935.7 respectively) as compared to the basic (no covariate) models 1-0 and 2-0. Models with host plant species ranked close to the selected model with a moderately inflated BIC relative to the selected model (Model 1-2: Δ_BIC_ = 971.1, Model 2-2: Δ_BIC_ = 933.2) and an important reduction in residual covariances relative to the basic model (compare Figures 2A and 2C). In comparison, including ecological variables alone deteriorated model fit with a greatly increased BIC (Model 1-3: Δ_BIC_ = 15088.8, Model 2-4: Δ_BIC_ = 11387.9) and a mild reduction of residual covariances (compare Figures 2A and 2B).

The selected model showed a good fit, with a tendency to overestimate low abundance values (Figure S3). As expected, model diagnostics revealed an excess of zeros and some over dispersion (Appendix S3) which PLN modeling is robust to (Chiquet *et al*. 2019). No temporal or spatial autocorrelation was detected in model residuals. The coefficients relative to host plant species, which represent the response of species abundances to host plants, had a strongly bimodal distribution (Figure 3A, right panel). For each fly, some plant species had very low coefficients (<-80), meaning a negligible effect of the corresponding host plant on fly abundances (approx. exp(−80) ≈ 1E-35) and we considered them as non-host plants. The other plants had much stronger coefficients (>-14) and were interpreted as host plants. Overall, this inferred realized host range was narrower than the laboratory-measured fundamental host range (Figure 3A, left panel), particularly for generalists. Only 10 fly-plant associations (out of 147) showed the reverse pattern (zero laboratory-measured fitness and yet strong inferred response to plant), suggesting marginal difficulties with measuring fitness in laboratory conditions. Among plants inferred as hosts from species abundance patterns (*i*.*e*., high coefficient values), coefficients correlated positively with fly laboratory-measured fitness for specialists but not for generalists (Figure 3B).

**Figure 3:**
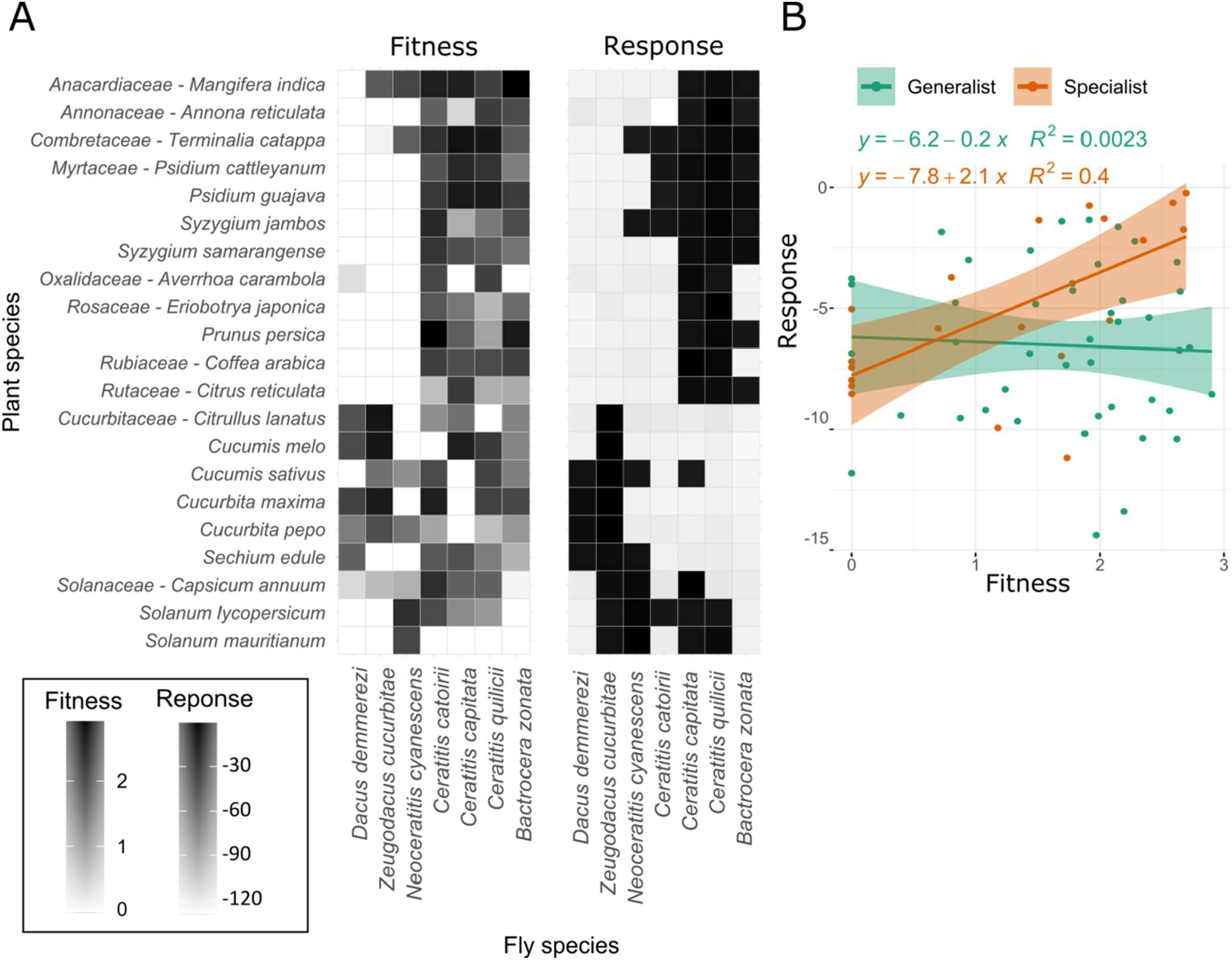
Comparison of fly species’ fundamental and realized host use. A) Fundamental host use as measured in the laboratory (left panel – fitness is the logarithm of the product preference and performance) and realized host use as inferred from regression coefficients relative to host plants in the fullest model (right panel – response is obtained from Model 2-5 on the seven-species dataset). B) Relationship between inferred responses to host plants and laboratory-measured fitness for specialists in orange and generalists in green on hosts detected as such in the field. Lines represent linear regressions with slope 95% confidence intervals as shadowed areas.

The coefficients relative to the first two axes of the FAMD on ecological covariates could be interpreted as responses of fly abundances to rainfall, temperature, and elevation (Figure 4). Rainfall had a low effect on the abundances of *B. zonata, C. capitata, C. quilicii*, and *D. demmerezi. Ceratitis catoirii, N. cyanenscens*, and *Z. cucurbitae* showed a propensity towards warm high-rainfall areas, while D. ciliatus seemed to prefer colder drier climates. *Ceratitis quilicii* and *C. catoirii* were not much affected by temperature. *Bactrocera zonata, C. capitata, D. ciliatus*, and *Z. cucurbitae* should thrive in low-elevation warm climates. *Dacus demmerezi* and to a lesser extent *N. cyanescens* seemed to prefer colder higher-elevation climates.

**Figure 4:**
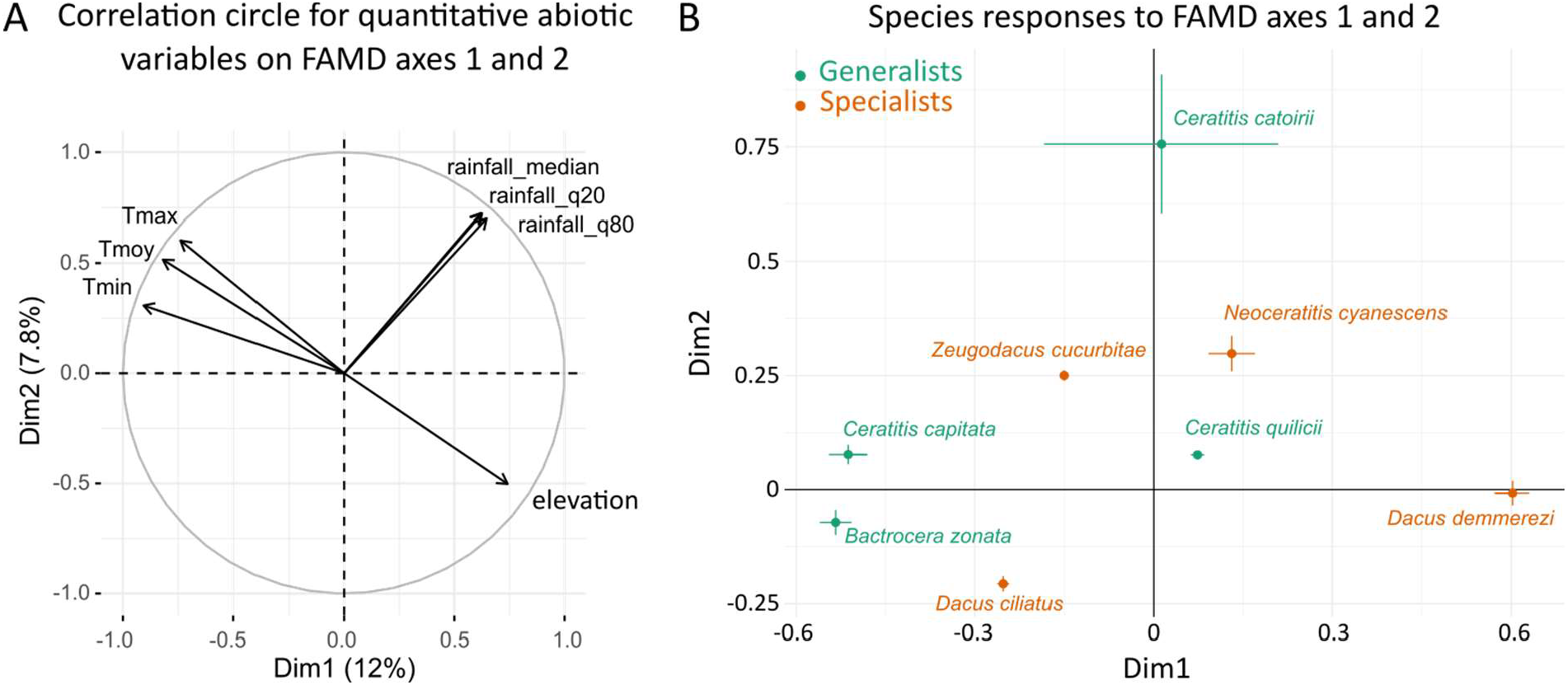
Species abundance responses to ecological variables. A) Correlation circle on the first and second axes of the FAMD (factorial analysis of mixed data) on ecological variables (Dim1 and Dim2, respectively). The first axis contrasts warm low-altitude sites and colder high-altitude sites. The second axis is a gradient of rainfall and maximal temperature (see Supplementary figure S3 for details on axes contributions). B) Regression slopes relative to Dim1 and Dim2, inferred under the fullest model on the 21 plant x 8 fly species dataset (Model 1-5). Error bars represent approximate confidence intervals (1.96 x standard errors). For the first axis (Dim1), negative slopes (*e*.*g*., *B. zonata* and *C. capitata*) can be interpreted as a positive effect of temperatures and a negative effect of elevation on species abundances. For the second axis (FAMD 2), positive slopes (*e*.*g*., *C. catoirii*) can be interpreted as a positive effect of rainfall on species abundances.

The selected model had full residual variance-covariance matrix on the eight-species dataset and diagonal residual variance-covariance matrix on the seven-species dataset. On the eight-species dataset its diagonal version ranked second with Δ_BIC_ = 428.9. Congruently, the residual variance-covariance matrix inferred under the selected model (Figure 2D) had very low covariance values for all pairs of fly species, all covariances being negative. This suggested possible, though weak, antagonist interactions. The largest residual correlations were observed between the three specialists D. demmerezi, D. ciliatus and Z. cucurbitae (residual correlations ranging from -0.031 to -0.112) and between the generalists B. zonata, C. quilicii and C. capitata (residual correlations from -0.025 to -0.090). Network inference, applied to the selected model, converged to one significant interaction between D. ciliatus and Z. cucurbitae, with a stability of 0.99 (*i*.*e*., detected in 99 out of 100 network inferences on bootstrapped data). Overall these results suggested that at the scale of the whole community competitive interactions between fly species only weakly affected their joint distributions and that fly species abundances were mainly explained by environmental covariates.

Considering models accounting for species fundamental niche (Table 1B), the best model with species traits included female preference, larval performance and ecological variables (Model 2-12, Δ_BIC_ = 5189.2). It ranked intermediate between the model with ecological variables alone and the best model. Such good performance of models with species traits suggests that the fundamental host range of fly species is an important determinant of fly species joint distributions. Whether in combination with ecological covariates or not, models with larval performance were slightly better than models with female preference.

### Cucurbitaceae

Focusing on Cucurbitaceae and the three fly species they hosted, *D. ciliatus, D. demmerezi* and *Z. cucurbitae*, the selected model was again the one including both host plant species and ecological covariates (Model 1-5, Table S5A). Its BIC was lower than the one of the basic model (Model 1-0, Δ_BIC_ = 1208.0). It assumed a full residual variance-covariance matrix and performed better than its diagonal version (Δ_BIC_ = 190.6). Network inference yielded two significant negative interactions between *D. ciliatus* and *Z. cucurbitae* (stability = 1.0) and between *D. demmerezi* and *Z. cucurbitae* (stability = 0.56). For all other models, versions with full residual matrix performed better than their diagonal counterparts. Besides, models including plant species alone or ecological covariates alone performed equivalently (Δ_BIC_ = 515.3 *vs*. Δ_BIC_ = 653.2). This is congruent with the idea that plant species affect fly distributions more similarly within the Cucurbitaceae family than at the scale of the 21-plant dataset, so that the influence of plant identity is less important for species distributions among Cucurbitaceae than on a wider set of plants.

For *D. demmerezi* and *Z. cucurbitae*, for which traits were available, models with species traits alone had poor performance, and displayed higher BIC than the basic model (Table S5B). However, the model with both species traits and ecological covariates ranked third just after the diagonal version of the selected model (Model 2-10, Δ_*BIC*_ = 124.9).

### Solanaceae

For Solanaceae and their associated fly species, *N. cyanescens, C. capitata*, and *C. quilicii*, the selected model included host plant species, ecological covariates and a diagonal residual variance-covariance matrix (Model 1-6, Table S6). It improved the BIC of the basic model by 201.8. It performed better than its full residual matrix version (Δ_BIC_ = 15.8) and congruently, applying network inference to the latter yielded no significant interaction between fly species. On this dataset, all models with a diagonal matrix performed better than their full residual matrix counterpart, further confirming the absence of detected interactions. Models with ecological covariates alone performed poorly (Model 1-3 Δ_BIC_ = 176.4 and Model 1-4 Δ_BIC_ = 162.0, respectively). Models with species traits were almost as good as their equivalent with host plant as a cofactor. For instance, the model with species traits and ecological covariates (Model 2-10) ranked second with Δ_BIC_ = 9.5. Female preference and larval performance performed equally well (Δ_BIC_ = 44.6 and Δ_BIC_ = 38.5 respectively) and almost as good as both traits together (Δ_BIC_ = 33.4), suggesting important correlation between the traits.

### Other plant families

The last dataset considers all families other than Cucurbitaceae or Solanaceae with *B. zonata, C. capitata, C. catoirii*, and *C. quilicii* (Table S7). The selected model included host plant species and ecological covariates (Model 1-5, Δ_BIC_ = 2639.7 with the basic model 1-0). It assumed a full residual variance-covariance matrix and performed slightly better than its diagonal version (Δ_BIC_ = 55.8). All models with a full residual matrix performed better than their diagonal counterpart. Network inference yielded one significant interaction between *B. zonata* and *C. quilicii* (stability = 1.0).

The model with only ecological covariates performed well (Model 1-3, Δ_BIC_ = 967.3), and almost as good as the model with host plant alone (Model 1-1, Δ_BIC_ = 684.1), suggesting redundancy between ecological information and plant identity. Models that included species traits without ecological covariates performed badly (Δ_BIC_ > 2220), and the model with species traits and ecological covariates (Model 2-10, Δ_BIC_ = 814.3) performed only slightly better than the model with ecological covariates only (Model 1-3, Δ_BIC_ = 967.3). Of the two traits, only female preference really improved model fit (Model 2-8 Δ_BIC_ = 888.9 *vs*. Model 2-12 Δ_BIC_ = 1050.0).

## Discussion

The determinants underlying the structure of a community of eight Tephritid fly species were deciphered. Modelling joint species abundances without accounting for any covariate (only intercepts and an offset) confirmed a major role of host use strategy on fly species abundances. Species abundances co-varied positively among generalists and among specialists and negatively between species of each of these groups. Common responses to environmental factors may cause positive residual correlations in species abundances, while divergent responses will imply negative correlations, potentially leading to incorrect interpretations of species interactions (Ovaskainen *et al*. 2016, Dormann *et al*. 2018). Accounting for environmental covariates strongly improved model fit and made all residual covariances almost completely vanish, particularly among groups, suggesting that no important environmental factor structuring the community has been missed. Obtaining long-term abundance data on all *a priori* relevant species of a given community is rarely possible. Here over the study period, one other fruit fly species (*Carpomya vesuviana*) was mentioned locally. It was considered very rare and only present on Indian jujube (*Ziziphus mauritiana*) in dry low-elevation areas (Quilici & Jeuffrault 2001, Franck *et al*. 2017). It was not detected in the 204 jujube samples of the full dataset. In addition, not all 108 plant species considered as potential fruit fly hosts were included. To evaluate the effect of omitted plants, we conducted the analysis on all plants with at least 10 samples with emerging flies (6434 samples, 36 plant species including the 21 studied in the laboratory) and obtained strikingly similar results (model ranking and estimates of regression coefficients, Appendix S3), comforting the idea that most relevant factors have been accounted for.

### Detection of competitive interactions

Some residual covariances remained non-negligible after accounting for fly species’ response to host plants between some generalists and between some specialists of Cucurbitaceae. They were all negative, suggesting a possible minor role of antagonistic interactions within specialists and within generalists. Only one of these residual covariations resisted the network inference process on the whole dataset (*D. ciliatus* - *Z. cucurbitae*, the two most abundant specialists of Cucurbitaceae). Two more significant covariations were detected when focusing on Cucurbitaceae (*Z. cucurbitae* - *D. demmerezi*) and on other plant families (*B. zonata* - *C. quilicii*) suggesting possible dependence of species interactions on host plants. On Cucurbitaceae, although qualitatively congruent with other independent empirical measures of host range (Vayssieres *et al*. 2008) and climatic niche (Vayssières & Carel 1999), host plant species and abiotic factors only moderately improved model fit. All three specialist flies found on Cucurbitaceae are able to thrive on any plant of this family (Charlery de la Masselière *et al*. 2017b) and competitive interactions between these fly species are highly plausible (Vayssieres *et al*. 2008). On other plant families, both host plants and abiotic factors clearly improved model fit, congruently with former interpretations of the system (Duyck *et al*. 2006a, Duyck *et al*. 2008). Responses of generalist species to abiotic factors were strikingly congruent with former independent laboratory experiments (Duyck & Quilici 2002, Duyck *et al*. 2004, Duyck *et al*. 2006a). There was redundancy between host plants and abiotic factors. Many of these plants are exploited but not planted (*e*.*g*., Myrtaceae). Their distributions are therefore more dependent on ecological factors than those of Solanaceae and Cucurbitaceae, which are mainly cultivated throughout the year in Réunion. On Solanaceae, no residual covariation was detected. Plant identity was the main determinant of species abundances, congruently with the idea that Solanaceae impose adaptive challenges on their fruit consumers through a variety of toxic compounds (Brévault *et al*. 2008), rendering host adaptation the main factor driving species abundances.

### A current competition ghost

Among-group covariation between specialists and generalists were mainly attributable to fly species’ adaptation to host use with a minor contribution of abiotic factors. Previous studies have highlighted differences in host adaptation between these fly species (Hafsi *et al*. 2016). Contrary to the specialists, which are mainly able to use their preferred hosts, the four generalists can thrive in numerous plant species, and have weak female preferences (Charlery de la Masselière *et al*. 2017b). Accordingly, specialists were seldom found in plants other than Cucurbitcaeae or Solanaceae. However, these results do not explain why generalists were so rarely found on Cucurbitaceae and Solanaceae. Competitive exclusion with specialists would be a natural hypothesis to explain this absence (Nakadai *et al*. 2018). Here, no competition among groups was detected. It is possible, however, that competition has already operated and that competitive exclusion has been so strong that generalists cannot be found on Cucurbitaceae. In PLN modelling, such absence could be interpreted as negative response of generalist abundances to Cucurbitaceae and be encapsulated in plant cofactor slopes rather than in residual covariances. It is precisely when competition is intense enough to cause niche partitioning that it can no longer be detected. This result evokes a well-known paradox in ecology termed ‘‘the ghost of competition past’’ (Connell 1980) according to which the observed differentiation in niches is the result of past interspecific competition.

To escape the paradox, knowledge about the fundamental niches of species through eco-evolutionary approaches could help settle whether species interactions are an important driver of species assemblages (Augustyn *et al*. 2016, Dormann *et al*. 2018). Laboratory measurements of larval performance and female preference on host plants were used in replacement of plant identity. Congruently with the community being essentially driven by host use, preference and performance clearly improved model fit. Interestingly, performance was more informative than preference, which was expected from previous knowledge that generalists’ preferences are uncorrelated to their performances (Charlery de la Masselière *et al*. 2017b). If competition truly shapes species abundances, and has not been detected, it is to be found in the difference between models with plant identity as a cofactor and models with species traits instead. Here we found a difference suggesting that competitive exclusion is at work. In terms of importance, from model rankings, host use patterns were the most important factor shaping species abundances, followed by abiotic factors and possibly a dose of competition.

This predominance of host plants as a structuring factor of phytophagous insect communities has been much debated, but congruent studies exist. In analyses of insect communities along road verges, Schaffers *et al*. (2008) found that the composition of plant communities was a much better predictor of insect and spider assemblages than environmental variables. Similarly Nakadai *et al*. (2018) found that sharing of host species was predominant among butterflies of the Japanese archipelago, suggesting that interspecific resource competition may not effectively determine community assembly patterns at regional scales. In an earlier review on the importance of competition in insects, Denno *et al*. (1995) pointed that only weak to moderate effect of competition should certainly be expected in phytophagous insects such as Tephritids due to their high mobility and weak aggregation behaviors. Experimental manipulations of competitive interactions in the field could offer a promising way to test the validity of the present inferences. These experiments would also be useful to unveil the role of other biotic interactions (*e*.*g*., natural enemies), as forces capable of modulating interspecific competition between fruit flies.

### Generalists vs. specialists

Overall specialists and generalists almost had very distinct realized host uses with different assembly rules. That specialists and generalists form separate interaction networks has already been highlighted, *e*.*g*., among soil microbial species (Barberan *et al*. 2012). It is well known that the predictability of assemblages differs between generalist and specialist phytophagous insects (Müller *et al*. 2011). This has led to the hypothesis that specialists would assemble according to the species-sorting paradigm of metacommunity ecology (Leibold *et al*. 2004), whereby species occurrences are mainly driven by habitat heterogeneity and local adaptation. Generalists’ assembly rules, on the other hand, would rather follow a mass-effect paradigm (Shmida & Wilson 1985), according to which sink populations, where the species is maladapted, can persist through a migration influx from source populations (Müller *et al*. 2011). Our results confirm this hypothesis. For specialists, a good agreement between the inferred host plant effects on species abundances and the laboratory measures of host adaptation suggested that specialists were mainly filtered by host plant characteristics. In contrast, generalists displayed no relationship between inferred and laboratory-measured host plant effects, suggesting that generalists were found on some hosts where their fitness is low and at low density on good hosts. Besides generalists use fruits whose availability is highly variable over time. Contrary to specialists, most of their hosts are not available all year long (Figure S4). This temporally variable habitat may trigger a dynamics of local extinction and recolonization, in which the roles of migration and stochasticity become more important than that of host adaptation and in which coexistence is possible despite fundamental niche overlap (Connell 1980, Chesson 2000).

## Data accessibility

Data and script are available online: http://doi.org/10.5281/zenodo.4569532

## Acknowledgements

This work is dedicated to Serge Quilici who has been leading prolific research on the Tephritids of Réunion his whole career. Cirad technicians Jim Payet and Serge Glénac made this study possible through their invaluable expertise with fly rearing and ecology. We also thank them as well as Antoine Franck, Christophe Simiand, and Patrick Turpin, for collecting field data over the years. Thomas Brequigny contributed to measuring larval traits during his internship. This work used images acquired within the framework of the CNES Kalideos device (Réunion site), which benefited from the “Programme Investissements d’Avenir” EQUIPEX of the French “Agence Nationale de la Recherche” on project GEOSUD bearing the reference ANR-10-EQPX-20. The images also required financial support the French Ministry of Agriculture and field data transmitted by the “Syndicat du Sucre de la Réunion” and the “SAFER de la Réunion”. BF, FC, FM, JC, MD, SR, and VR received the financial support of the French “Agence Nationale de la Recherche” project NGB (ANR-17-CE32-011). EF, FC, PFD, and VR were funded by the European Union (European Regional Development Fund, ERDF contract GURDT I2016-1731-0006632), the Conseil Régional de La Réunion, and the Centre de Coopération Internationale en Recherche Agronomique pour le Développement (CIRAD). AH was funded by the “Ministère de l’Enseignement supérieur et de la Recherche Scientifique de la Tunisie”. This study used the facilities provided by the Plant Protection Platform (3P, IBISA), Saint-Pierre, Réunion, France.

Version 4 of this preprint has been peer-reviewed and recommended by Peer Community In Ecology (https://doi.org/10.24072/pci.ecology.100080).

## Supplementary material

### Appendix S1: Additional materials & methods

#### Details on species table

Among the 12872 initial samples, only samples that fulfilled to following conditions were kept: (i) GPS coordinates could be retrieved (n=9715 in 41 plant species), (ii) at least one Tephritid fly emerged (n=6455 in 41 plant species), (iii) from plant species which had been successfully sampled at least 10 times in the dataset (n=6434 in 36 plants), and (iv) from plant species that were also characterized in the laboratory (n=4918 in 21 plants). In the resulting dataset, 97351 individual flies were counted. Samples covered 104 field sessions all year round over the 1991-2009 period (Tables S1 and S2). The number of flies per sample varied from 1 to 2244. Samples differed in terms of fruit number and weight. The number of flies varied from 1 to 188 per individual fruit and from 1.1E-3 to 15.8 flies per gram of fruit.

**Table S1:** Number of samples (*n*) by year in the full 21-plant dataset

| Year | 1991 | 1992 | 1993 | 1994 | 1995 | 1996 | 1997 | 1998 | 2001 | 2002 | 2003 | 2004 | 2005 | 2008 | 2009 |
| --- | --- | --- | --- | --- | --- | --- | --- | --- | --- | --- | --- | --- | --- | --- | --- |
| $n$ | 4 | 15 | 31 | 52 | 80 | 38 | 434 | 1143 | 83 | 523 | 634 | 19 | 205 | 191 | 1466 |

**Table S2:**
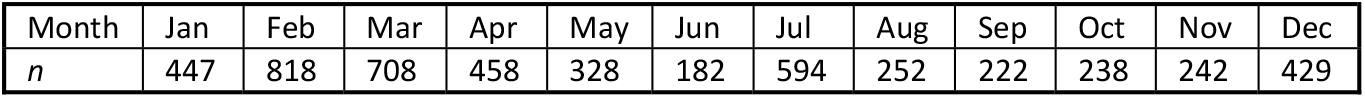
Number of samples (*n*) by month in the full 21-plant dataset

| Month | Jan | Feb | Mar | Apr | May | Jun | Jul | Aug | Sep | Oct | Nov | Dec |
| --- | --- | --- | --- | --- | --- | --- | --- | --- | --- | --- | --- | --- |
| $n$ | 447 | 818 | 708 | 458 | 328 | 182 | 594 | 252 | 222 | 238 | 242 | 429 |

The number of samples per plant species (Table S3) varied from 12 (*Averrhoa carambola*) to 1105 (*Cucurbita maxima*). The most represented plant families were the Cucurbitaceae (six plant species and 2374 samples), the Myrtaceae (four plant species and 1102 samples) and the Combretaceae (one plant species with 853 samples).

**Table S3:**
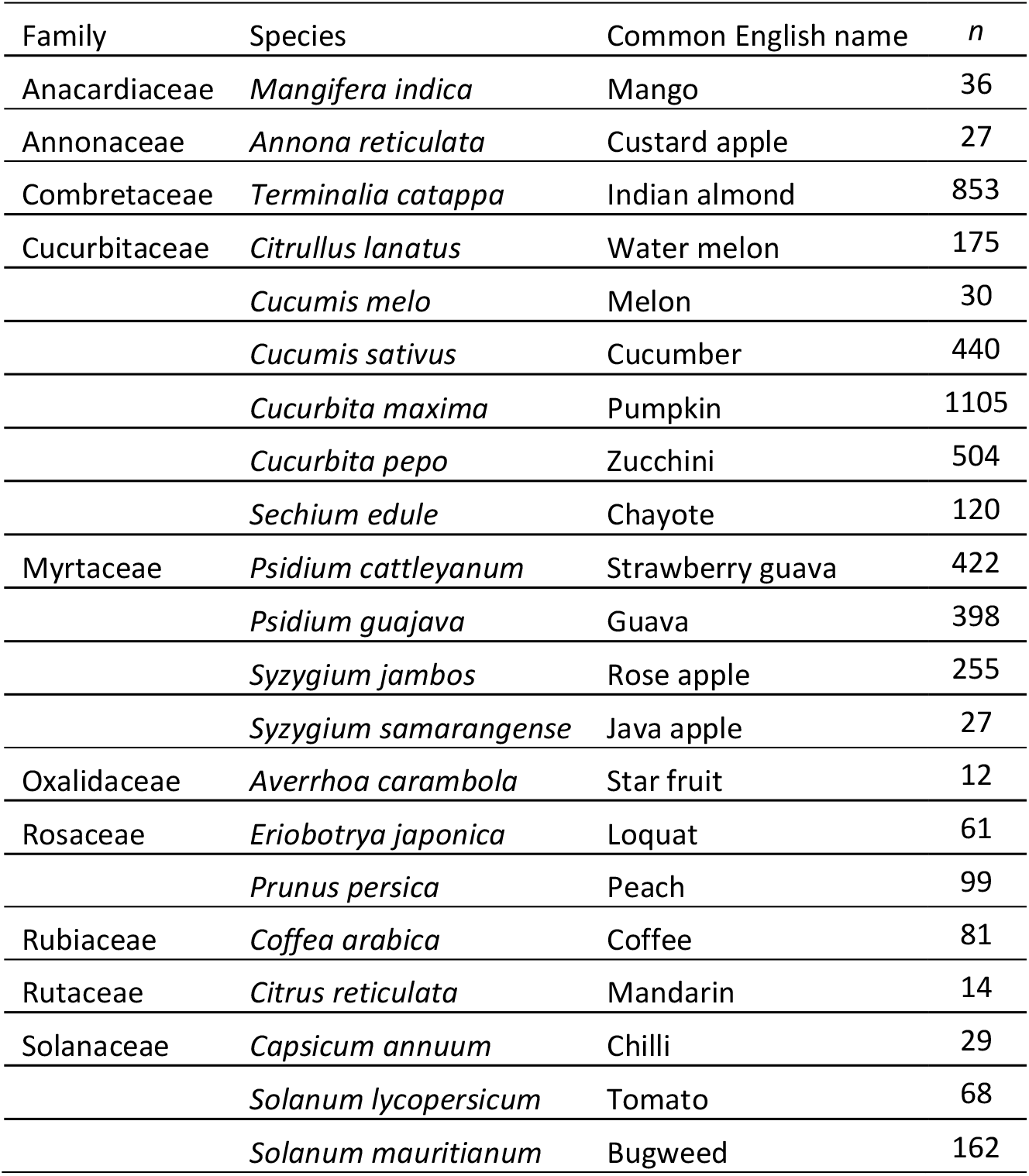
Number of samples (*n*) for each of the 21 plant species retained in the study

Plant species differed in terms of the patterns of occurrence of the eight fly species (Figure 2B). In particular, Cucurbitaceae were mostly used by the three species known as specialists of this plant family, *D. ciliatus* (14047 flies in 1264 samples), *D. demmerezi* (6335 flies in 276 samples) and *Z. cucurbitae* (21830 flies in 1463 samples). Other fly species were detected in fewer than 10 Cucurbitaceae samples. Solanaceae mainly hosted the specialist species *N. cyanescens* (1205 flies in 225 samples). They were also, to a lesser extent, hosts of *C. capitata* (116 flies in 32 samples), and *C. quilicii* (131 flies in 22 samples). Other fly species were detected in less than 5 Solanaceae samples. The other plant species (12 plant species) were mainly used by the four generalist species, *i*.*e*., *B. zonata* (11517 flies in 728 samples), *C. capitata* (8735 flies in 537 samples), *C. quilicii* (33149 flies in 1583 samples), *C. catoirii* (291 flies in 65 samples). Other fly species were detected in less than 5 samples.

Co-occurrences between fly species were frequent. 1148 samples out of 4918 (23.3%) contained more than one species, and 759 out of 4335 samples with a single fruit (17%) contained more than one fly species. Co-occurrences between fly species were therefore much structured by whether the considered plant species belonged to Cucurbitaceae, Solanaceae or other families (Table S4).

**Table S4:** Species co-occurences in the full 21-plant dataset (*n* = 4918 samples)

|  | *Bactrocera zonata* | *Ceratitidis quilicii* | *Ceratitidis capitata* | *Ceratitidis catoirii* | *Neoceratitidis cyanescens* | *Dacus ciliatus* | *Dacus demmerezii* | *Zeugodacus cucurbitae* |
| --- | --- | --- | --- | --- | --- | --- | --- | --- |
| *Bactrocera zonata* | 728 | 198 | 120 | 1 | 0 | 0 | 0 | 0 |
| *Ceratitidis quilicii* | 198 | 1605 | 309 | 57 | 12 | 1 | 0 | 1 |
| *Ceratitidis capitata* | 120 | 309 | 570 | 31 | 15 | 1 | 0 | 2 |
| *Ceratitidis catoirii* | 1 | 57 | 31 | 66 | 1 | 1 | 0 | 1 |
| *Neoceratitidis cyanescens* | 0 | 12 | 15 | 1 | 237 | 3 | 1 | 5 |
| *Dacus ciliatus* | 0 | 1 | 1 | 1 | 3 | 1266 | 107 | 454 |
| *Dacus demmerezii* | 0 | 0 | 0 | 0 | 1 | 107 | 276 | 133 |
| *Zeugodacus cucurbitae* | 0 | 1 | 2 | 1 | 5 | 454 | 133 | 1467 |

#### Ecological covariables

Elevation (in meters) of each sample was obtained from the Digital Elevation Model Litto3D® co-produced by the French IGN (National Geographic Institute) and the SHOM (Marine Oceanographic Hydrographic Service). Pluviometry was obtained from layers produced by M. Mezino from 143 CIRAD and Meteo-France meteorological stations, containing isohyets of (i) the minimal rainfall observed in the 20% most humid years of the 1986-2016 period (ii) the minimal rainfall observed in the 20% driest years of the 1986-2016 period and (iii) the median annual rainfall over the 1986-2016 period. For each sample, the value of closest isohyet was retained for each of the three variables. Temperature was characterized by the (i) minimal, (ii) mean and (iii) maximal annual temperature over the 1987-2017 period as interpolated by M. Mezino from 73 CIRAD and Meteo-France meteorological stations raw data. Land use around each sample location was obtained from a 12-categories layer produced by supervised classification of Pléiades 2018 images (Dupuy and Gaetano, 2019 doi:10.18167/DVN1/WKAJZO, CIRAD Dataverse, V1).

#### Additional details on FAMD on ecological covariables

**Figure S1:**
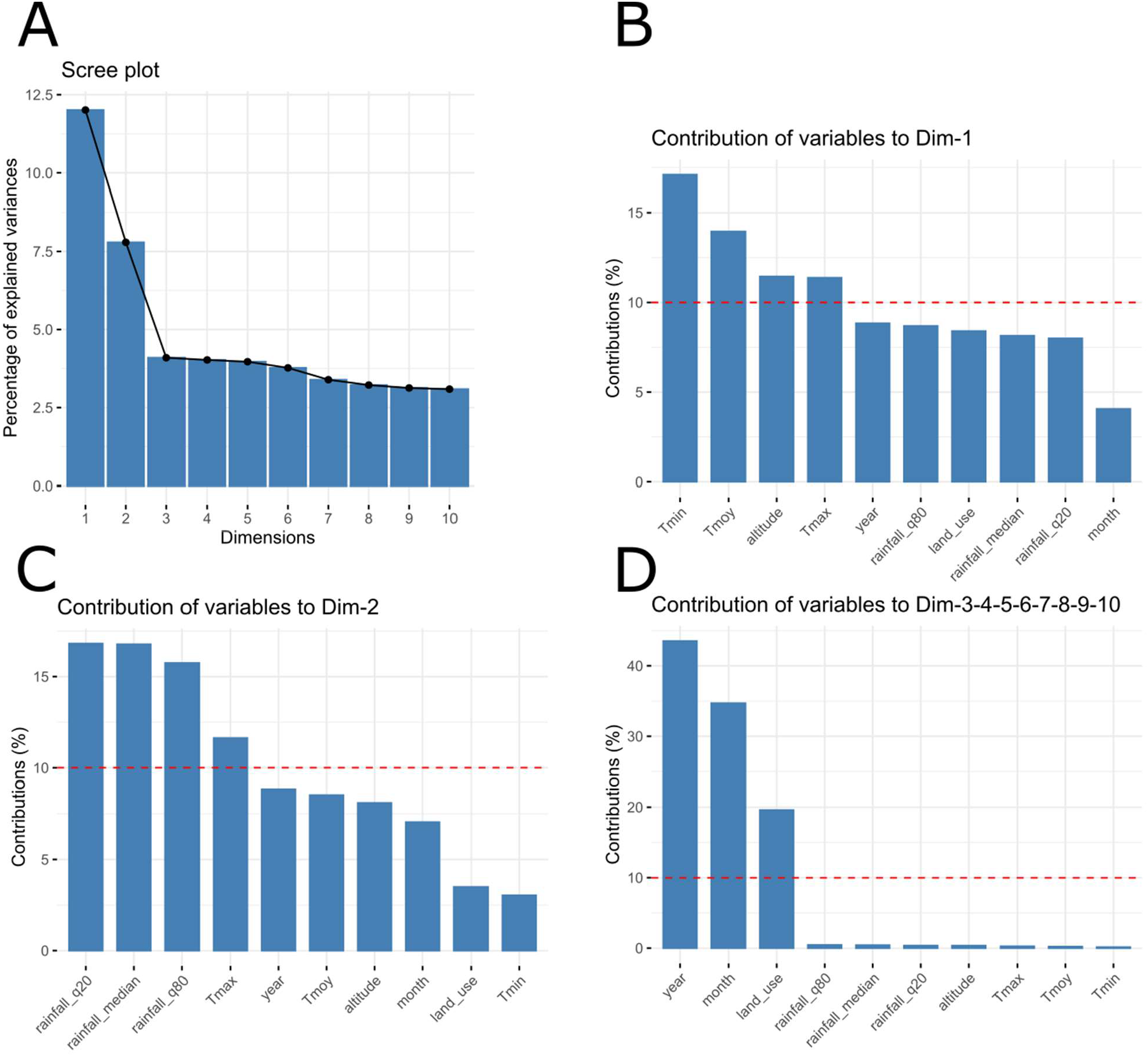
Interpretation of the axes of the FAMD on ecological covariables. A) Percentage of variance explained by each of the 10 axes of the FAMD. The first and second FAMD axes explain 19.8 % of the variance. C-D) Contribution of individual ecological covariates to Axis 1, 2 and other axes, respectively. Axis 1 is mainly contributed by temperature and elevation. Axis 2 is mainly contributed by rainfall and maximal temperature. Other axes are contributed by qualitative variables (year, month and land use).

### Appendix S2: Complementary results

#### Model selection on the three sub-datasets

##### Cucurbitaceae

**Table S5:** Model selection on the Cucurbitaceae dataset (*n* = 2347). Models are ranked by increasing BIC (from best to worst). *k* is the number of parameters. *L* is the log-likelihood. Δ_BIC_ is the BIC difference between any focal model and the best one.

| Model | Covariates | Residual matrix | $k$ | $L$ | BIC | $\Delta_{\text{BIC}}$ |
| --- | --- | --- | --- | --- | --- | --- |
| A) Model set 1 (6 plants x 3 flies) |  |  |  |  |  |  |
| Model 1-5 | Plant + Eco | Full | 54 | -14168.3 | 28756.3 | 0.0 |
| Model 1-6 | Plant + Eco | Diagonal | 51 | -14275.3 | 28947.0 | 190.6 |
| Model 1-1 | Plant | Full | 24 | -14542.5 | 29271.6 | 515.3 |
| Model 1-3 | Eco | Full | 39 | -14553.2 | 29409.5 | 653.2 |
| Model 1-2 | Plant | Diagonal | 21 | -14661.3 | 29485.7 | 729.4 |
| Model 1-4 | Eco | Diagonal | 36 | -14649.7 | 29579.2 | 822.8 |
| Model 1-0 | None | Full | 9 | -14947.2 | 29964.3 | 1208.0 |
| B) Model set 2 (6 plants x 2 flies) |  |  |  |  |  |  |
| Model 2-5 | Plant + Eco | Full | 35 | -7311.6 | 14881.6 | 0.0 |
| Model 2-6 | Plant + Eco | Diagonal | 34 | -7361.7 | 14974.4 | 92.8 |
| Model 2-10 | Preference + Performance + Eco | Diagonal | 28 | -7399.9 | 15006.5 | 124.9 |
| Model 2-3 | Eco | Full | 25 | -7419.8 | 15024.2 | 142.6 |
| Model 2-8 | Preference + Eco | Diagonal | 26 | -7421.5 | 15035.1 | 153.5 |
| Model 2-12 | Performance + Eco | Diagonal | 26 | -7423.9 | 15039.8 | 158.2 |
| Model 2-4 | Eco | Diagonal | 24 | -7461.8 | 15100.9 | 219.3 |
| Model 2-1 | Plant | Full | 15 | -7570.5 | 15251.8 | 370.2 |
| Model 2-2 | Plant | Diagonal | 14 | -7654.1 | 15411.5 | 529.9 |
| Model 2-0 | None | Full | 5 | -7722.3 | 15481.6 | 600.0 |
| Model 2-7 | Preference | Diagonal | 6 | -7739.0 | 15522.4 | 640.8 |
| Model 2-9 | Preference + Performance | Diagonal | 8 | -7745.6 | 15550.4 | 668.7 |
| Model 2-11 | Performance | Diagonal | 6 | -7801.2 | 15646.7 | 765.1 |

##### Solanaceae

**Table S6:** Model selection on the Solanaceae dataset (*n* = 259). Models are ranked by increasing BIC (from best to worst). *k* is the number of parameters. *L* is the log-likelihood. Δ_BIC_ is the BIC difference between any focal model and the best one.

| Model | Covariates | Residual matrix | $k$ | $L$ | BIC | $\Delta_{\text{BIC}}$ |
| --- | --- | --- | --- | --- | --- | --- |
| Model 1-6 | Plant + Eco | Diagonal | 42 | -824.2 | 1881.7 | 0.0 |
| Model 2-10 | Preference + Performance + Eco | Diagonal | 42 | -829.0 | 1891.2 | 9.5 |
| Model 1-5 | Plant + Eco | Full | 45 | -823.8 | 1897.5 | 15.8 |
| Model 2-12 | Performance + Eco | Diagonal | 39 | -842.6 | 1901.7 | 20.0 |
| Model 2-8 | Preference + Eco | Diagonal | 39 | -846.5 | 1909.5 | 27.8 |
| Model 1-2 | Plant | Diagonal | 12 | -922.3 | 1911.2 | 29.4 |
| Model 2-9 | Preference + Performance | Diagonal | 12 | -924.2 | 1915.1 | 33.4 |
| Model 2-11 | Performance | Diagonal | 9 | -935.1 | 1920.2 | 38.5 |
| Model 1-1 | Plant | Full | 15 | -921.1 | 1925.6 | 43.8 |
| Model 2-7 | Preference | Diagonal | 9 | -938.2 | 1926.3 | 44.6 |
| Model 1-4 | Eco | Diagonal | 36 | -921.9 | 2043.8 | 162.0 |
| Model 1-3 | Eco | Full | 39 | -920.8 | 2058.1 | 176.4 |
| Model 1-0 | None | Full | 9 | -1016.8 | 2083.5 | 201.8 |

##### Other plant families

**Table S7:** Model selection on the dataset with other plant families. Models are ranked by increasing BIC (from best to worst). *k* is the number of parameters. *L* is the log-likelihood. Δ_BIC_ is the BIC difference between any focal model and the best one.

| Model | Covariates | Residual matrix | $k$ | $L$ | BIC | $\Delta_{\text{BIC}}$ |
| --- | --- | --- | --- | --- | --- | --- |
| Model 1-5 | Plant + Eco | Full | 98 | -12218.5 | 25195.0 | 0.0 |
| Model 1-6 | Plant + Eco | Diagonal | 92 | -12269.6 | 25250.8 | 55.8 |
| Model 1-1 | Plant | Full | 58 | -12715.3 | 25879.1 | 684.1 |
| Model 1-2 | Plant | Diagonal | 52 | -12796.6 | 25995.3 | 800.3 |
| Model 2-10 | Preference + Performance + Eco | Diagonal | 56 | -12788.1 | 26009.3 | 814.3 |
| Model 2-8 | Preference + Eco | Diagonal | 52 | -12840.9 | 26083.9 | 888.9 |
| Model 1-3 | Eco | Full | 54 | -12872.4 | 26162.4 | 967.3 |
| Model 2-12 | Performance + Eco | Diagonal | 52 | -12921.4 | 26245.1 | 1050.0 |
| Model 1-4 | Eco | Diagonal | 48 | -12968.4 | 26308.1 | 1113.1 |
| Model 2-9 | Preference + Performance | Diagonal | 16 | -13650.6 | 27424.9 | 2229.8 |
| Model 2-7 | Preference | Diagonal | 12 | -13855.9 | 27804.7 | 2609.7 |
| Model 1-0 | None | Full | 14 | -13863.2 | 27834.7 | 2639.7 |
| Model 2-11 | Performance | Diagonal | 12 | -13905.2 | 27903.1 | 2708.1 |

#### Host plants availability

**Figure S2:**
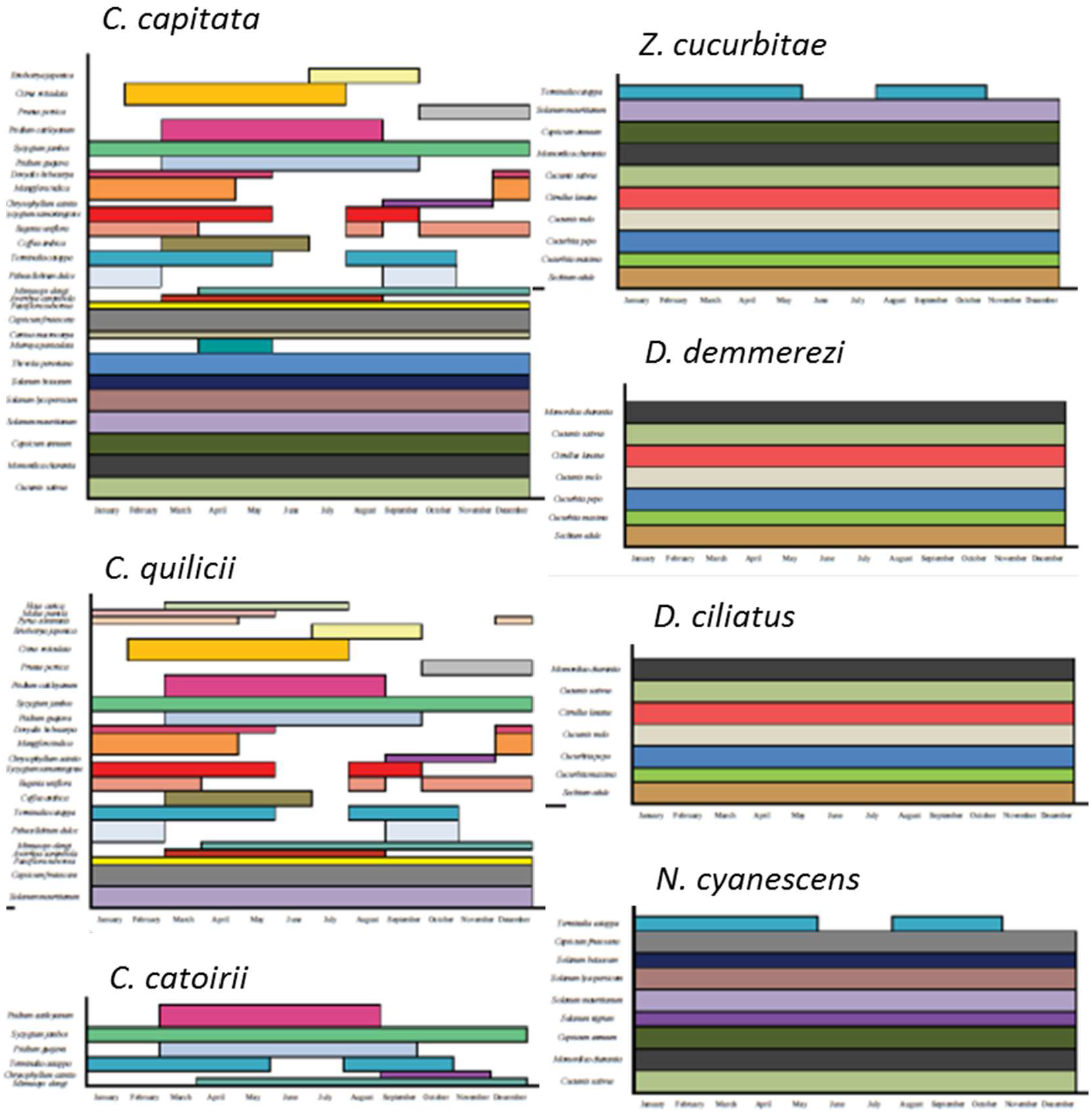
Availability of host plants along the year for seven fly species from Charlery de la Masselière *et al*. (2017a, Supplementary material Figure S1). Only major host plants are reported. Specialist species tend to rely on hosts with year-long availability, while generalists also have host with short fructification periods in their diet.

### Appendix S3: Stability and robustness of results

#### Model diagnostics

Model diagnostics were conducted using the R package DHARMa (Hartig, 2020; Figure S3 below).

**Figure S3.**
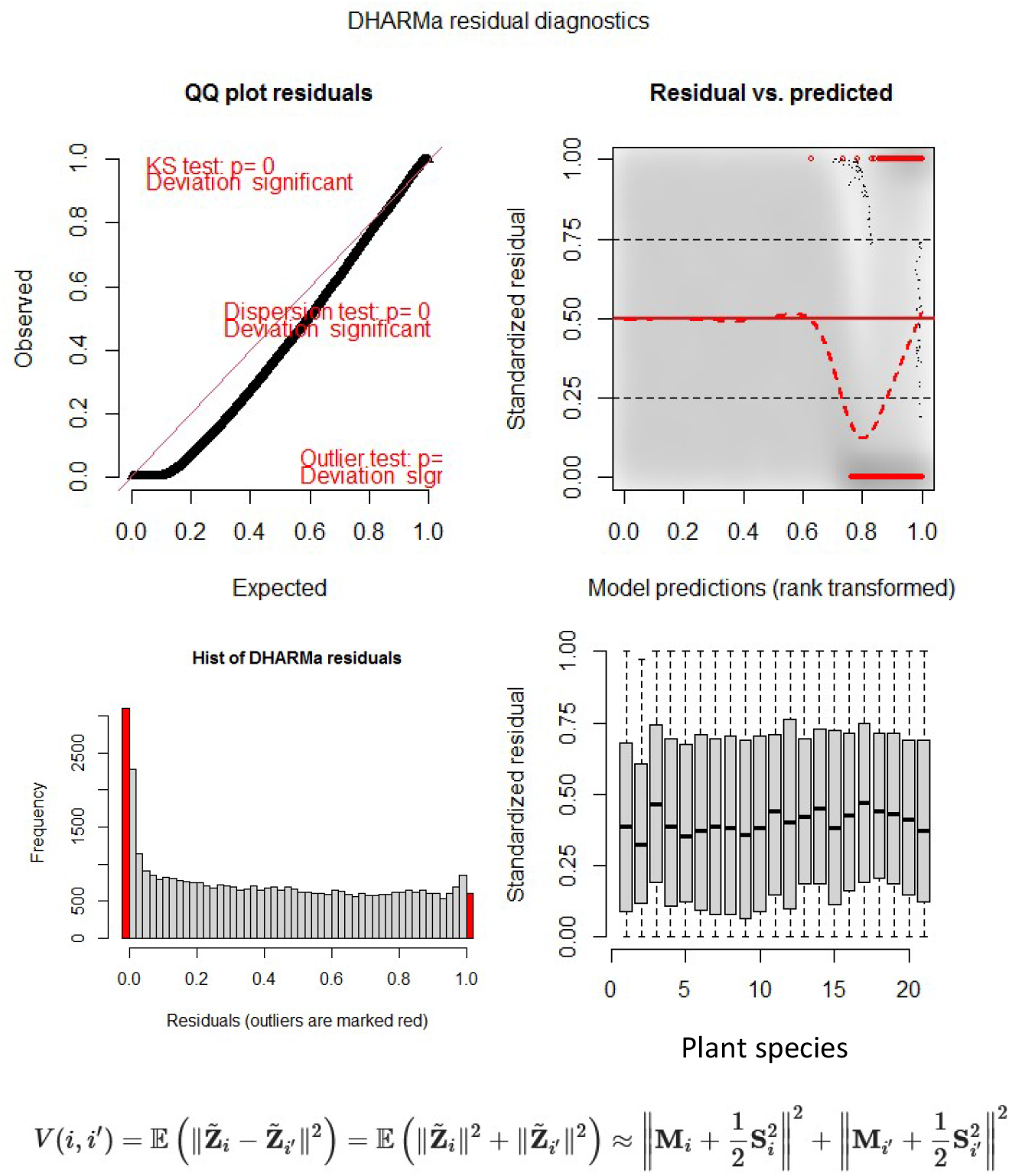

Even on the most complete model, an excess of zeros was detected likely in relation with fly species host ranges: for any given fly species not all 21 studied plants are compatible hosts, therefore introducing many zeros in the complete 8-fly dataset. No particular tendency of residuals on predictors was identified (plant shown here, FAMD axes not shown).

To further explore potential spatial or temporal trends in model residuals, we studied the variogram of residuals, *i*.*e*., how the estimated variance of the difference between two sites of this quantity changes as a function of spatial distance of distance in sampling date (equation below, Figure S4):

Correlations between the variance of residuals and spatial distance, difference in sampling month or difference in year of sampling were weak (0.0433, -0.0049, and 0.0211, respectively).

**Figure S4:**
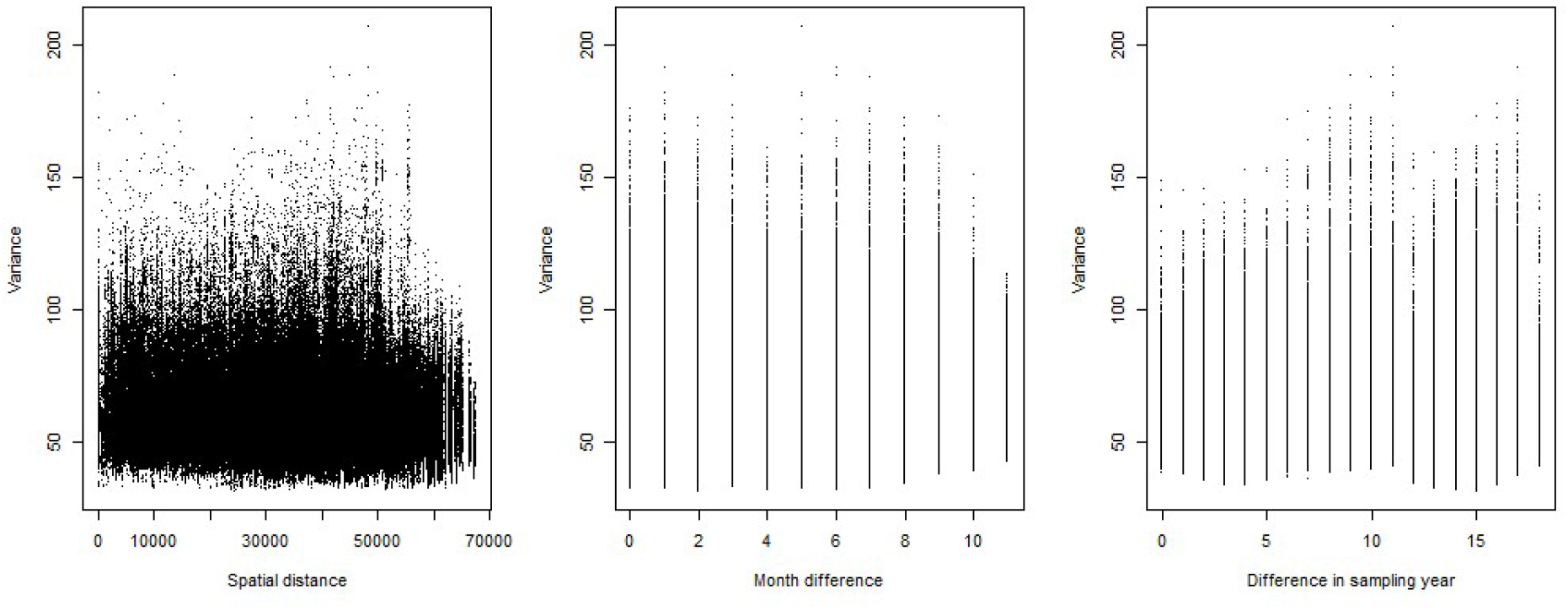
Variogram of the residuals of the fullest model on the full 21 plant x 8 fly species dataset (Model 1-5) with respect to spatial distance, difference in the month of sampling or difference in year of sampling.

Lastly, we explored the relationship between observed and fitted species abundances (log scale) in the best models on the full 21 plant x 8 fly species dataset (Figure S5, Model 1-5 on the left panel and model 1-6 on the right panel). Data were relatively well fitted except that both models tended to overestimate low abundance values.

**Figure S5.**
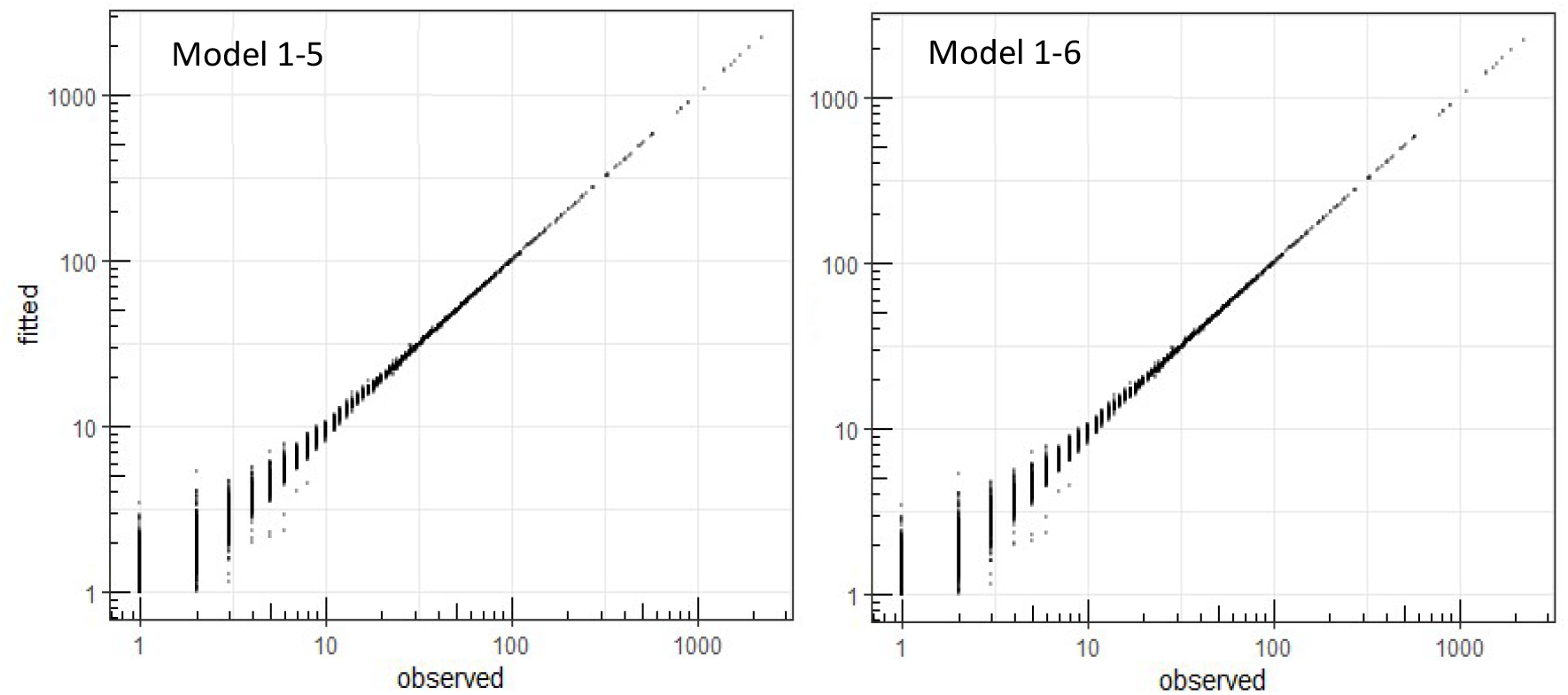

#### BIC robustness in PLN models comparisons

**Figure S6.**
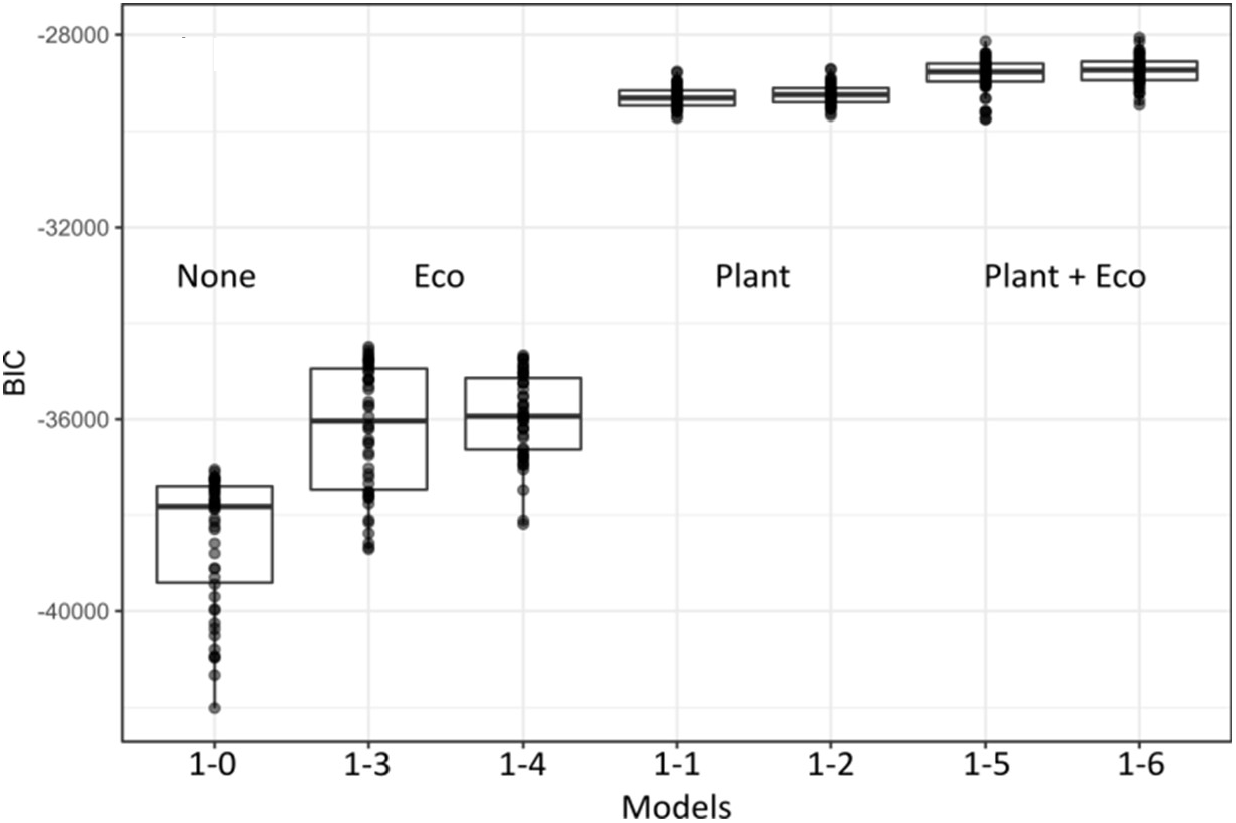

To estimate the magnitude of the uncertainty associated to model selection, we have proceeded to simulations by bootstrapping samples 50 times and conducting model comparison on the bootstrapped datasets for all models of Table 1 (no covariate, plant, ecological factors, both with either full or diagonal residual matrix on the full 21 plant x 8 fly species dataset).

#### Stability of sparse structure in residual matrix

**Figure S7.**
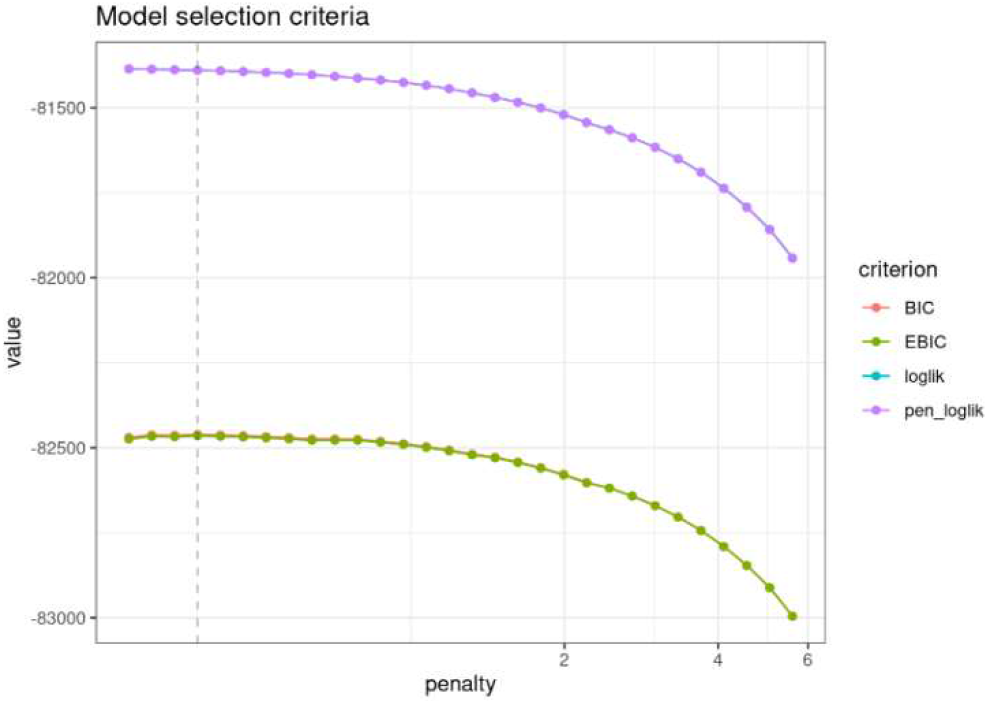

**Figure S8.**
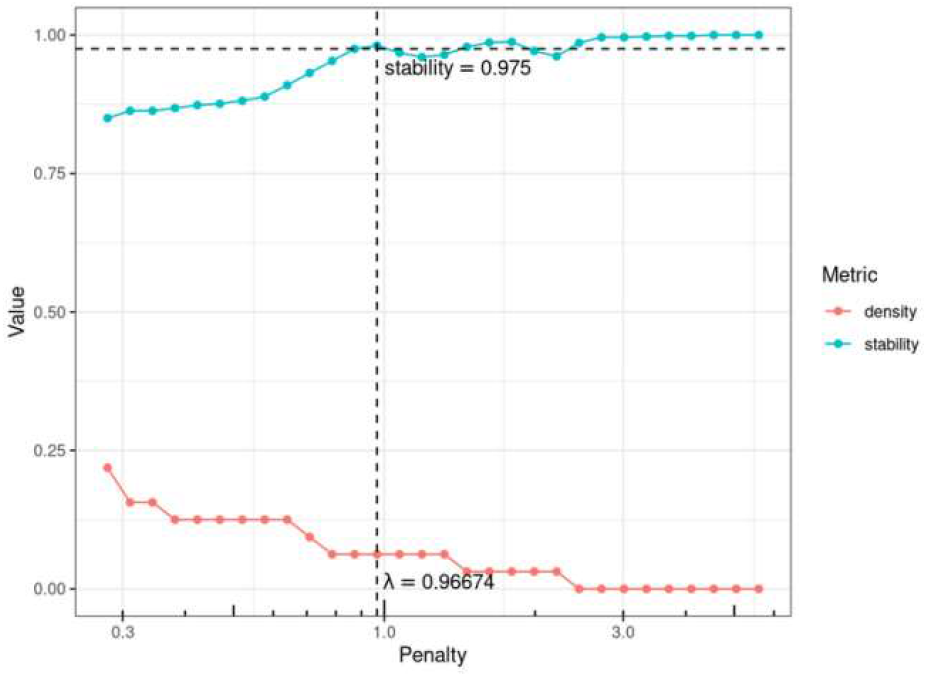

The function PLNnetwork() allows digging into the residual variance-covariance matrix and infer which species pairs still have with significant unexplained covariation in abundance after accounting for covariates. In this approach, the residual matrix is formalized as a network, which nonzero edges point to species pairs with significantly unexplained covariations. The function builds a series of 40 models with varying penalties over the number of edges in the resulting network (*i*.*e*., nonzero residual covariances), and allows comparing them using model selection criteria (Figure S7). The stability of network edges, can be evaluated using StARS (Liu *et al*. 2010), which performs resampling to evaluate the robustness of the network along the path of solutions. It allows selecting networks with varying stability criteria (Figures S8 and S9).

**Figure S9:**
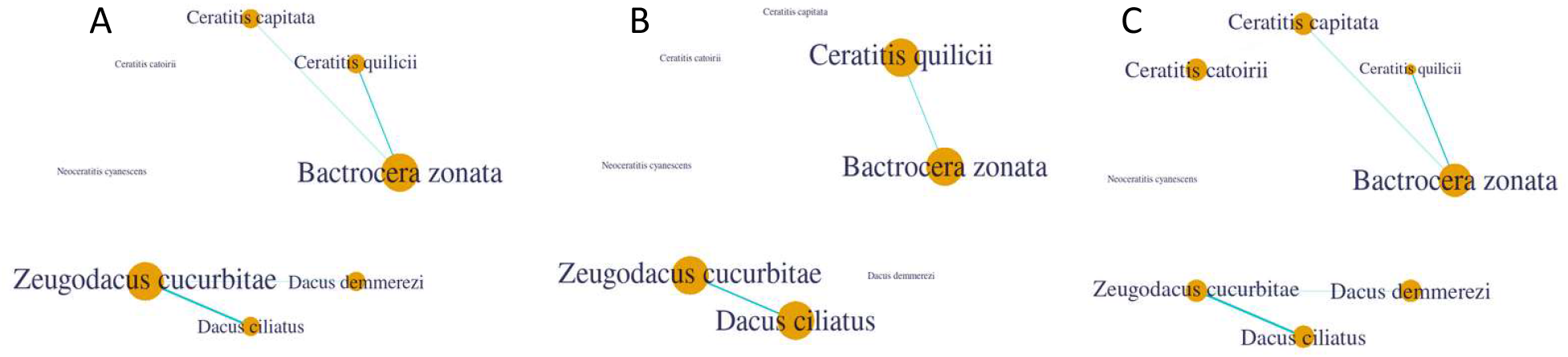
Various networks inferred. A. Network with the best BIC criterion. B. Network with a stability of 0.95. C. Network with a stability of 0.75. Whatever the model selection criterion, significant residual associations are found between the generalists *B. zonata* and *C. quilicii*, and between the specialists *Z. cucurbitae* and *D. ciliatius*.

#### Inferred realized host use using 36 host plants

The full initial dataset contained 12872 from 40 host plants. Keeping only samples with GPS coordinates, at least one individual fly and at least 10 samples per host plant led to a filtered dataset with 6434 samples and 36 host plants, including the 21 ones further studied in the laboratory. Conducting PLN modelling on the 36-plant datatset led to the following results.

**Figure S10:**
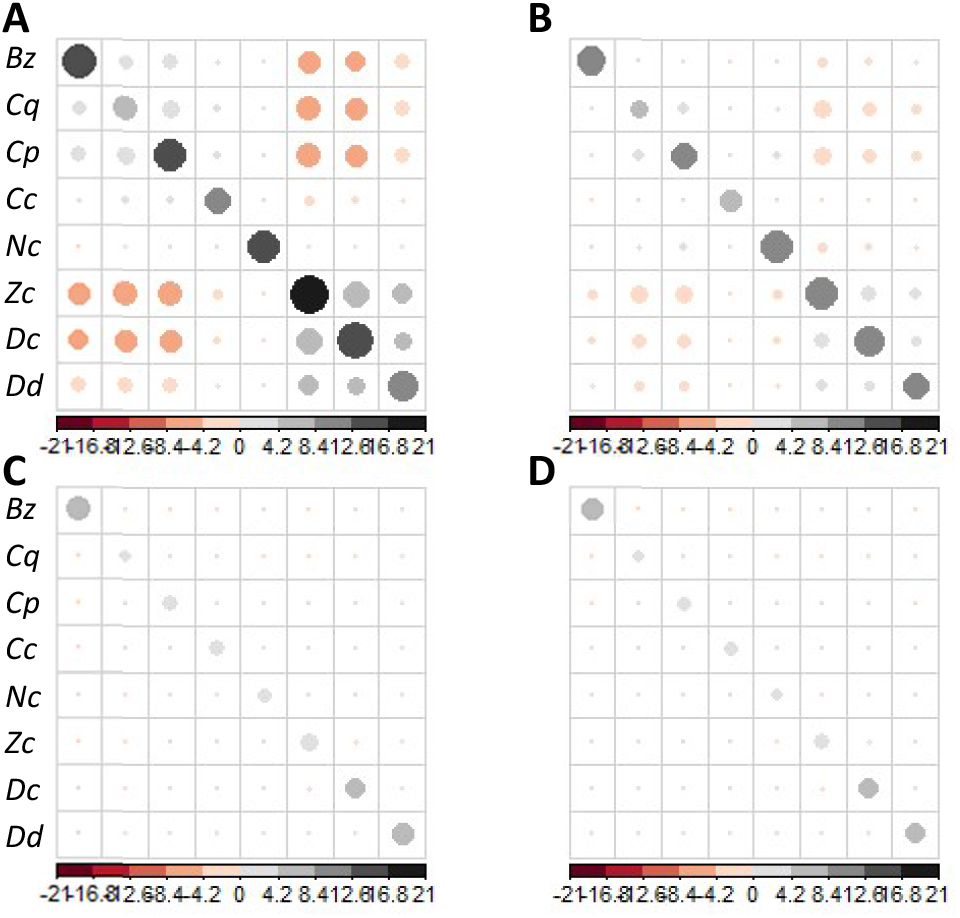
Residual variance-covariance matrices obtained after PLN model fitting on species abundances. A) Without any covariate (Model 1-0). B) With ecological covariates (Model 1-3). C) With plant species as a cofactor (Model 1-1). D) **B** With both plant species and ecological covariates (Model 1-5).

**Table S8:** Model selection on the extended dataset (*n* = 6434). Models are ranked by increasing BIC (from best to worst). *k* is the number of parameters. *L* is the log-likelihood. Δ _BIC_ is the BIC difference between any focal model and the best one.

| Model | Covariates | Residual Matrix | $k$ | $L$ | BIC | $\Delta_{\text{BIC}}$ |
| --- | --- | --- | --- | --- | --- | --- |
| Model 1-6 | Plant + Eco | Diagonal | 376 | -35018.92 | 73335.11 | 0.00 |
| Model 1-5 | Plant + Eco | Full | 404 | -34955.09 | 73452.99 | 117.87 |
| Model 1-2 | Plant | Diagonal | 296 | -35933.11 | 74461.95 | 1126.83 |
| Model 1-1 | Plant | Full | 324 | -35869.94 | 74581.15 | 1246.04 |
| Model 1-4 | Eco | Diagonal | 96 | -47094.34 | 95030.54 | 21695.42 |
| Model 1-3 | Eco | Full | 124 | -48796.13 | 98679.67 | 25344.55 |
| Model 1-0 | None | Full | 44 | -50013.67 | 100413.20 | 27078.08 |

**Figure S11:**
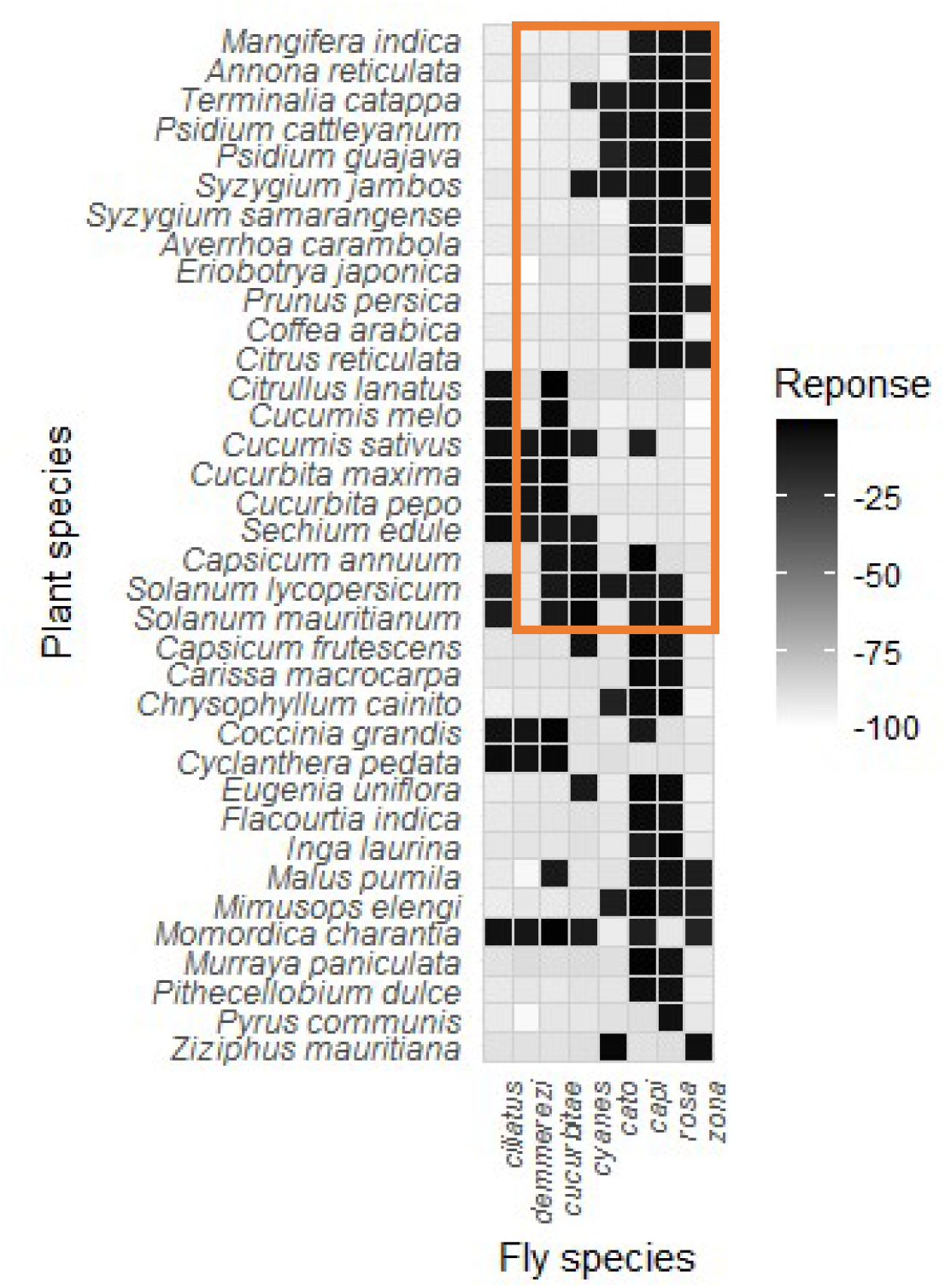
Realized host use as inferred from regression coefficients relative to host plants in the fullest model (Model 1-5 on the eight-species dataset). The orange rectangle contains plant-fly associations studied in the main text (Figure 3).

**Figure S12:**
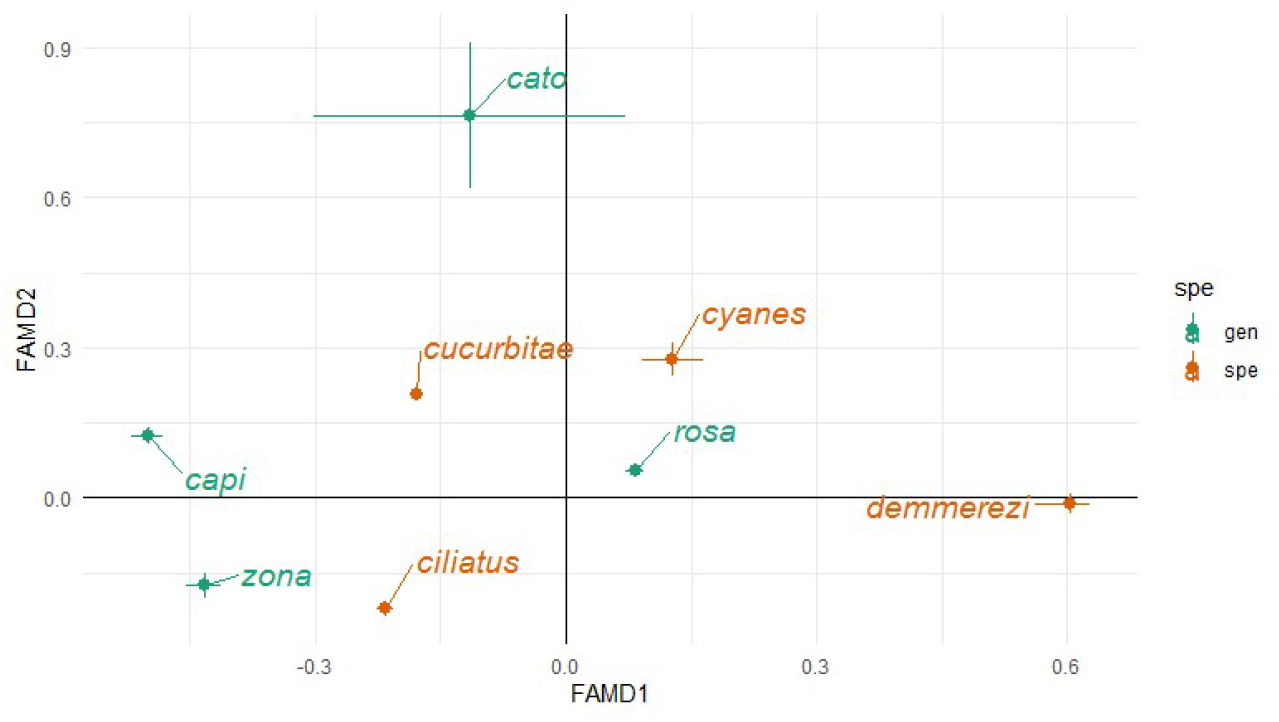
Species abundance responses to ecological variables inferred through regression coefficients relative to Dim1 and Dim2, inferred under the fullest model on the 36 plant x 8 fly species dataset (Model 1-5). Error bars represent approximate confidence intervals (1.96 x standard errors).

